# Diverse somatic genomic alterations in single neurons in chronic traumatic encephalopathy

**DOI:** 10.1101/2025.03.03.641217

**Authors:** Guanlan Dong, Chanthia C. Ma, Shulin Mao, Samuel M. Naik, Katherine Sun-Mi Brown, Gannon A. McDonough, Junho Kim, Samantha L. Kirkham, Jonathan D. Cherry, Madeline Uretsky, Elizabeth Spurlock, Ann C. McKee, August Yue Huang, Michael B. Miller, Eunjung Alice Lee, Christopher A. Walsh

**Affiliations:** Division of Genetics and Genomics, Manton Center for Orphan Disease Research, Boston Children’s Hospital; Boston, MA, USA; Department of Pediatrics, Harvard Medical School; Boston, MA, USA; Bioinformatics and Integrative Genomics Program, Harvard Medical School; Boston, MA, USA; Harvard-MIT MD-PhD Program, Harvard Medical School; Boston, MA, USA; Program in Biological and Biomedical Sciences, Harvard Medical School; Boston, MA, USA; Division of Neuropathology, Department of Pathology, Brigham and Women’s Hospital, Harvard Medical School; Boston, MA, USA; Department of Biological Sciences, Sungkyunkwan University; Suwon, South Korea; Veterans Affairs (VA) Boston Healthcare System, US Department of Veteran Affairs; Boston, MA, USA; Alzheimer’s Disease Research Center and Chronic Traumatic Encephalopathy Center, Chobanian and Avedisian School of Medicine, Boston University; Boston, MA, USA; Department of Pathology and Laboratory Medicine, Chobanian and Avedisian School of Medicine, Boston University; Boston, MA, USA; Department of Neurology, Chobanian and Avedisian School of Medicine, Boston University; Boston, MA, USA; Broad Institute of MIT and Harvard; Cambridge, MA, USA; Howard Hughes Medical Institute; Boston, MA, USA

## Abstract

Chronic traumatic encephalopathy (CTE) is a neurodegenerative disease that is linked to exposure to repetitive head impacts (RHI), yet little is known about its pathogenesis. Applying two single-cell whole-genome sequencing methods to hundreds of neurons from prefrontal cortex of 15 individuals with CTE, and 4 with RHI without CTE, revealed increased somatic single-nucleotide variants in CTE, resembling a pattern previously reported in Alzheimer’s disease (AD). Furthermore, we discovered remarkably high burdens of somatic small insertions and deletions in a subset of CTE individuals, resembling a known pattern, ID4, also found in AD. Our results suggest that neurons in CTE experience stereotyped mutational processes shared with AD; the absence of similar changes in RHI neurons without CTE suggests that CTE involves mechanisms beyond RHI alone.

## Main Text

Chronic traumatic encephalopathy (CTE) develops years after exposure to repetitive head impacts (RHI) and is most often found in contact sport athletes, such as boxers and American football players (*1–5*). CTE is diagnosed by postmortem neuropathological examination, based on the presence of a pathognomonic lesion consisting of perivascular hyperphosphorylated tau neurofibrillary tangles in neurons at the depths of the cortical sulci (*1, 2*). While tau deposition is common to CTE and Alzheimer’s disease (AD) (*4, 6*), CTE is characterized by distinct neuropathological features, tau molecular structural conformation (*7, 8*), and clinical symptoms (*5, 9*). The precise mechanisms by which RHI drives tau neurodegeneration are poorly understood. Somatic mutations are well-known drivers of cellular proliferation in neoplasia (*10–12*), but also accumulate in non-neoplastic cells and have been observed to increase in neurons with age (*13–17*). Multiple studies using single-cell whole-genome sequencing (scWGS) (*17, 18*) and single-molecule duplex sequencing (*16, 19, 20*) have converged on accumulation rates of 16-17 somatic single-nucleotide variants (SNVs) per year and 2-3 somatic small insertions and deletions (Indels) per year in neurons from neurotypical controls. Furthermore, we previously reported a significantly higher burden of distinct mutational signatures in neurons from individuals with AD and other neurodegenerative disorders (*18, 21*). To further investigate the role of somatic mutations in neurodegeneration, we applied scWGS to the amplified DNA from single neuronal nuclei isolated from CTE, RHI without CTE, AD, and neurotypical control individuals (Fig. 1A), using both strand-agnostic and strand-specific amplification methods to assess somatic mutations in CTE.

**Fig. 1.**
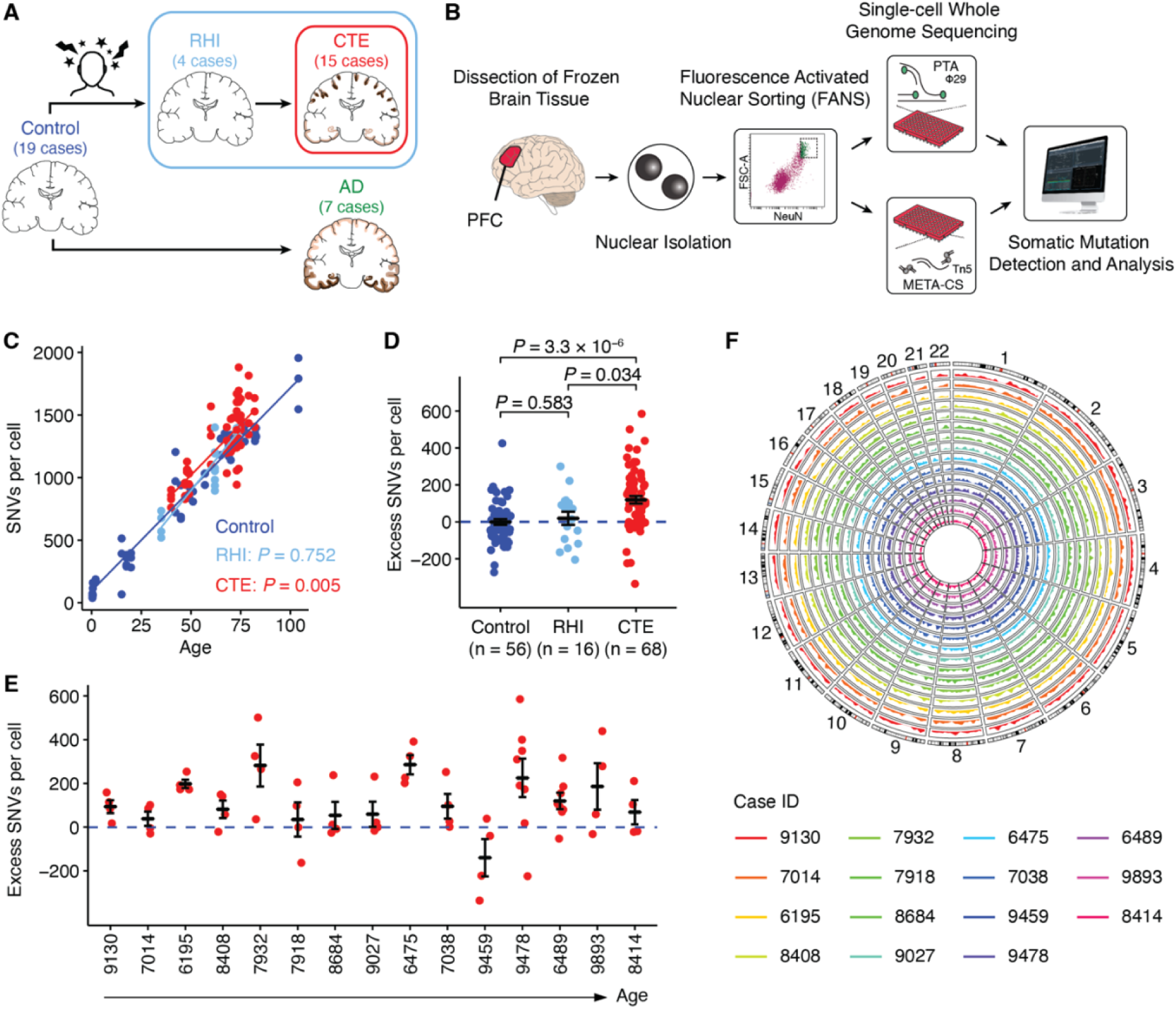
Study design and somatic SNV burden in CTE neurons. (**A**) Cohort design: neurotypical control, repetitive head impacts (RHI), chronic traumatic encephalopathy (CTE), and Alzheimer’s disease (AD) postmortem human brain tissue. Illustrations of CTE and AD show characteristic tau pathology patterns (in brown). The number of cases included in this study are shown for each clinical condition. (**B**) Single-cell whole-genome sequencing (WGS) experimental approach. Nuclei are isolated from postmortem human brain tissue and subjected to fluorescence-activated nuclear sorting for the neuronal nuclear marker NeuN. Nuclei are sorted one-per-well, lysed, and subjected to primary template amplification (PTA) and multiplexed end-tagging amplification of complementary strands (META-CS). Amplified genomic DNA is then assayed using WGS to identify somatic mutations. PTA data is used to determine somatic mutation burden, while META-CS data is used to identify strand-related signatures. (**C**) Increased somatic SNV burden in CTE (red) brains compared to RHI (light blue) brains and neurotypical controls (dark blue). Somatic SNV burden (from each neuron as a point) estimated by SCAN2 is fitted against age by clinical conditions using a LME model (neurotypical control: dark blue; RHI: light blue, *P* = 0.752; CTE: red, *P* = 0.005). (**D**) CTE neurons show a significant excess of somatic SNV burden (*P* = 3.3 × 10^-6^, two-tailed Wilcoxon test) while RHI neurons show no significant difference compared to neurotypical controls (*P* = 0.583, two-tailed Wilcoxon test). (**E**) Excess somatic SNVs in each CTE case ordered by increasing age. The dashed blue line shows SNVs attributable to age (zero excess). (**F**) Circos plot showing the PTA somatic SNV density distribution of CTE cases across the whole genome. Each CTE case is depicted by color in a circular track. Cases are ordered by increasing age same as (E) from the inner track to the outer track.

## Results

### Somatic SNVs in CTE neurons

We acquired frozen brain tissue of the dorsolateral prefrontal cortex (PFC) from individuals with a history of RHI exposure, with and without CTE, as well as neurotypical controls. We isolated nuclei of single neurons by staining for the nuclear stain DAPI, the pan-neuronal marker NeuN, and sorted neurons using fluorescence-activated nuclei sorting (FANS), gating specifically for the largest NeuN-positive nuclei (Fig. 1B, fig. S1). This gating method isolates pyramidal excitatory neurons with a purity of greater than 99% (*18*). We sorted single nuclei into individual wells of 96-well plates, then performed whole-genome amplification using primary template amplification (PTA) (*22*) or multiplexed end-tagging amplification of complementary strands (META-CS) (*19*). The amplified DNA underwent multiple screening and quality control steps, ensuring that only well-amplified genomes were used for sequencing. In total, using PTA, we sequenced the genomes of 68 neurons from 15 cases of CTE (exposed to RHI with neuropathologically verified CTE), 16 neurons from 4 cases of RHI (exposed to RHI without CTE), and compared these with 56 neurons from 19 neurotypical controls (*23*) and 28 neurons from 7 cases of AD (*18*) (Table 1, tables S1-2).

**Table 1.**
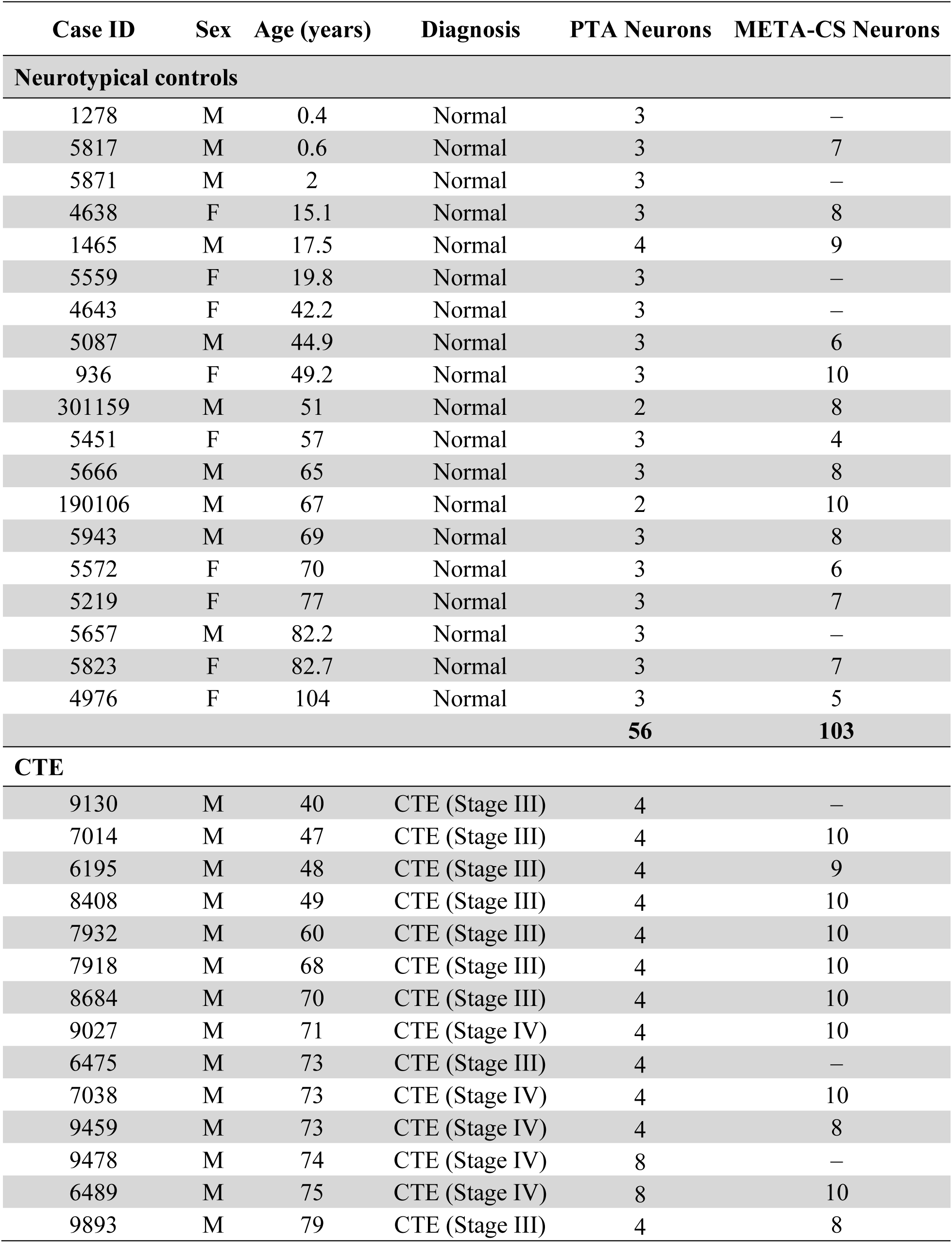
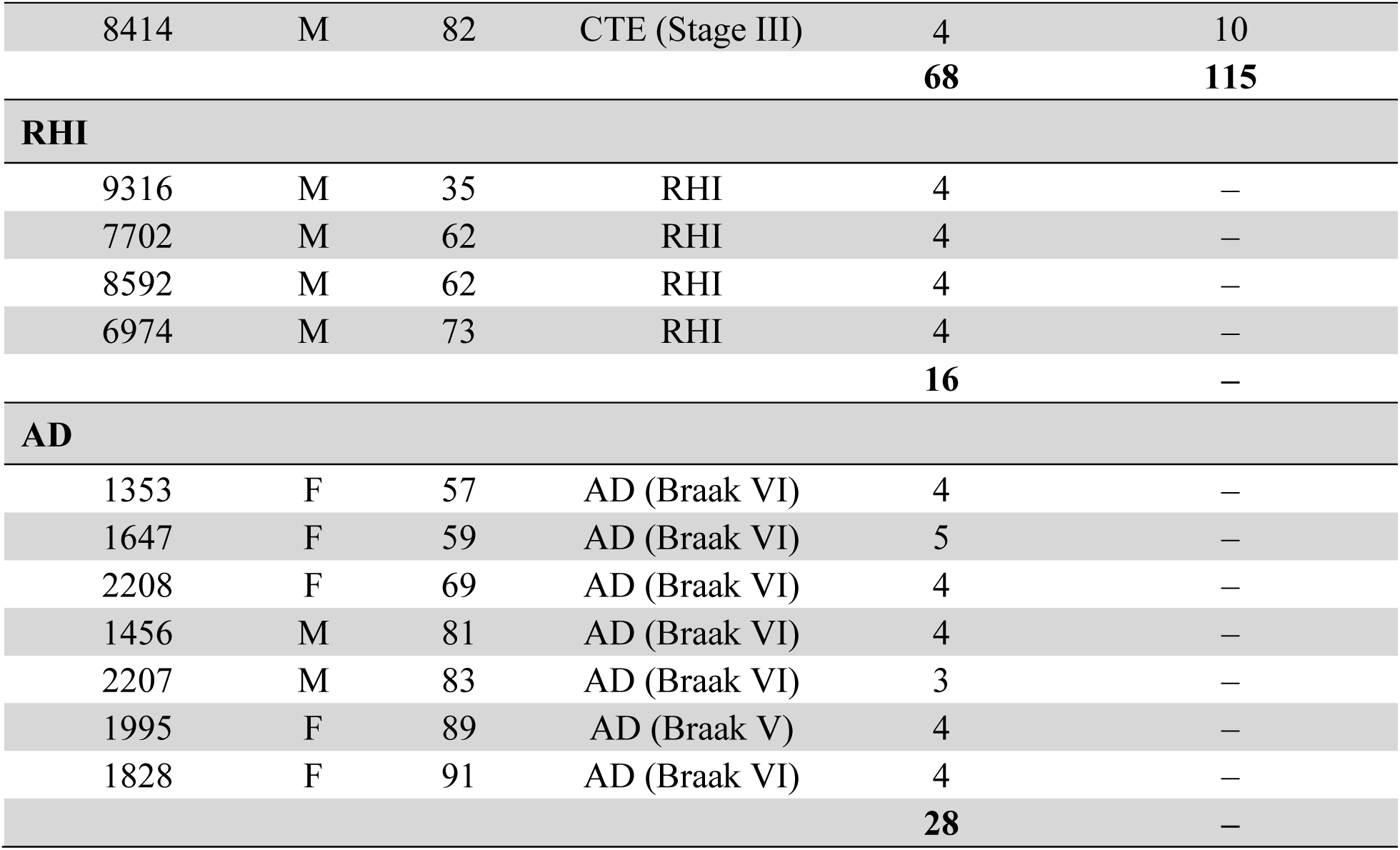
Sample information with number of neurons analyzed using two scWGS methods.

We identified both somatic SNVs and Indels in each single PTA neuron using SCAN2 (*17*) and found a significant increase of somatic SNVs in CTE, averaging 111 more SNVs per cell compared to controls (*P* = 0.005, linear mixed-effects (LME) model; Fig. 1C). In contrast, somatic SNV burdens in the RHI group were similar to controls (*P* = 0.752, LME model; Fig. 1C). These results suggest that increased somatic SNVs in CTE arise not from RHI exposure alone, but from additional factors associated with the development of CTE and the elevated DNA damage. All CTE and RHI brain samples were collected and processed at theUNITE Brain Bank to control for potential batch effects. After adjusting for age, CTE neurons also showed a significant excess of somatic SNVs compared to control neurons (*P* = 3.3 × 10^-6^, two-tailed Wilcoxon test; Fig. 1D). The increased somatic SNV burden in CTE remained significant after controlling for additional covariates representing sample and sequencing quality (see Methods, fig. S2). Furthermore, while neurons in each case showed some variation, we observed greater mean somatic SNVs in nearly all CTE cases than the burden attributable to normal aging (Fig. 1E).

We observed that somatic mutations were broadly distributed across each neuron’s genome (Fig. 1F). As CTE pathology is localized primarily at the depths of the cortical sulci relative to gyri (*1*), we explored the relationship between cortex topology and somatic mutations, but found no significant difference in somatic SNV burden between sulcal neurons and gyral neurons in CTE (*P* = 0.43, two-tailed Wilcoxon test; fig. S3). Previous studies reported a significant dose-response relationship between years of playing American football and CTE pathology, risk, and severity (*6, 24–26*). However, we observed no significant association between somatic SNV burden and years of playing football within our CTE samples after controlling for age (*P* = 0.235, LME model), which implies that the duration of RHI exposure does not directly lead to increased somatic SNV burden, in line with the similar somatic burden between RHI and controls.

To investigate the nature of excess somatic SNVs in CTE, we used a modified duplex META-CS method (*19*) (see Methods) to profile a total of 115 neurons from 12 CTE cases and 103 neurons from 14 neurotypical control cases with paired PTA scWGS data (Table 1). We distinguished variants representing double-stranded mutations and single-stranded DNA lesions, respectively (fig. S4A). Signature decomposition on somatic SNVs from the PTA-profiled neuron using META-CS-derived strand-oriented signatures showed on average 87% double-stranded signature contribution (fig. S4B), indicating that PTA-identified somatic SNVs are predominantly double-stranded, albeit not exclusively. This aligns with the observation that neurotypical controls show a highly similar rate of age-related SNV increase when profiled by either PTA (*17*) or strand-specific methods (*16, 19*).

### Mutational signatures of somatic SNVs in CTE neurons

To investigate potential sources of somatic SNVs in CTE neurons, we performed mutational signature analysis on the PTA scWGS data from CTE and neurotypical control neurons. First, we decomposed the somatic SNVs into Signatures A and C, which were previously identified in single human neurons (*21*) (Fig. 2A, B). Signature A closely resembles SBS5 from the COSMIC database (v.3.2) (*27*), a clock-like signature that is associated with aging. Signature C resembles SBS8, a signature associated with deficient transcription-coupled nucleotide excision repair (TC-NER) (*21, 28*), along with other single base substitution (SBS) signatures, which has also been linked with oxidative damage in AD neurons (*18*). The contribution of Signature A increased with age without showing a significant difference between CTE and control neurons (*P* = 0.088, LME model; Fig. 2A). In contrast, we observed a significantly larger contribution of Signature C in CTE neurons, with an average excess of 60 SNVs per cell (*P* = 7.7 × 10^-4^, LME model; Fig. 2B). We then compared the signature contributions to 28 previously reported AD neurons (*18*), in which AD showed a greater contribution of both Signature A and Signature C compared to age-matched controls (*P* = 0.009 and 0.007, LME model; fig. S5). To better understand the age- and disease-associated contributions of the two signatures, we calculated the ratio of each signature’s total SNVs to age-contributed SNVs. We observed a commensurate contribution of Signature A (ratio ∼ 1) across CTE, AD, and controls, and an elevated contribution of Signature C (ratio > 1) in CTE and AD (Fig. 2C), suggesting shared mutational mechanisms in both neurodegenerative tauopathies. Notably, when we compared the relative contribution of the two signatures after adjusting for age, we found that Signature C accounted for a greater proportion of the increase in CTE than in AD (Fig. 2D). META-CS data identified a mean contribution of Signature C ranging from 24% to 26% in double-stranded SNVs (dsSNVs) and from 67% to 69% in single-stranded SNVs (ssSNVs) for both CTE and age-matched controls (fig. S6A). This finding, combined with the predominant contribution of dsSNVs (∼87%) in PTA neurons, suggests that a substantial amount of Signature C is double-stranded. We did not observe a significant difference in the relative contribution of Signature C between CTE and age-matched controls for either dsSNVs or ssSNVs (*P* = 0.702 and 0.219, two-tailed Wilcoxon test; fig. S6B). While the precise mechanism of Signature C and SBS8-like variants has not been definitively determined, the C>A mutations in this pattern are associated with alterations in reactive oxygen species (ROS) in other contexts (*29*) and represent one possible contributor to disease pathogenesis.

**Fig. 2.**
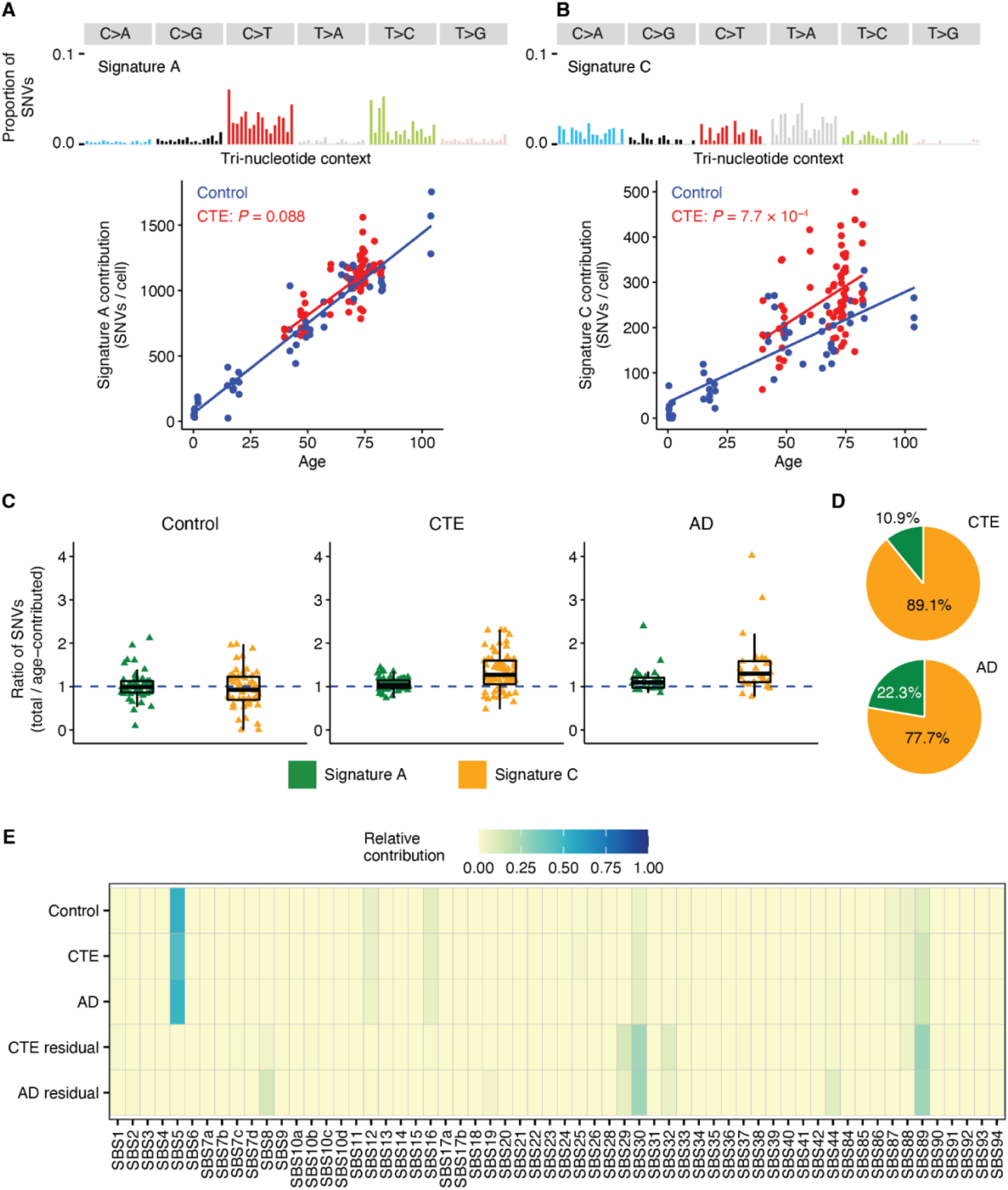
Somatic SNV mutational signatures in CTE neurons. (**A, B**) Somatic SNV burden separated by contributions from Signatures A (A) and C (B) in CTE (red) and neurotypical controls (dark blue) neurons. These signatures decompose somatic SNV burden into age- and disease-specific effects. Their contributions are fitted against age by clinical conditions using a LME model (A, Signature A, CTE: *P* = 0.088; B, Signature C, CTE: *P* = 7.7 × 10^-4^). (**C**) Ratios of each signature’s total contribution to age-related contribution in neurotypical control, CTE, and AD neurons. Age-contributed SNVs of each signature are obtained from the LME model in neurotypical controls. A ratio > 1 indicates a higher contribution from the signature compared to age-matched controls. Bars in each box plot from top to bottom show the first, second (median), and third quartile; whiskers extend 1.5 interquartile range (IQR). (**D**) Relative contribution of Signatures A and C in CTE and AD after adjusting for age. Based on the ratios shown in (C), the relative contribution of each signature is calculated by removing the median ratio of neurotypical controls (representing age effect) from the median ratio of each disease. (**E**) Mutational spectra of neurotypical control, CTE, and AD neurons are fitted to the COSMIC SBS database of cancer mutational signatures. Residual mutational patterns for CTE and AD are obtained by subtracting mutation profiles of age-matched controls from those of CTE and AD to show disease-specific contributions.

In addition to mapping the somatic SNVs to previously characterized signatures, we conducted *de novo* signature analysis using non-negative matrix factorization (NMF) (*30*). After determining the optimal number of signatures (fig. S7A), we identified two *de novo* signatures, N1 and N2 (fig. S7B). While N1 and N2 shared many similarities with Signatures A and C (fig. S7C), there were also substantial overlaps between N1 and N2 (e.g., SBS5 and SBS16) which may limit their usability for interpreting somatic SNV accumulation, as they may not represent distinct biological processes. Therefore, to delve deeper into other factors that may contribute to the excess somatic SNVs in CTE, we aggregated mutations based on their disease status (neurotypical control, CTE, and AD) and fitted them to COSMIC SBS signatures (Fig. 2E). The aging effect represented by SBS5 was dominant across all three groups. However, once we subtracted the pattern of age-matched controls from CTE and AD, we were able to obtain residual patterns that were more specific to each disease (fig. S7D). Interestingly, we found an elevated contribution of SBS29 and SBS32 in CTE that is almost absent in controls. SBS29 is a signature associated with tobacco chewing, yet distinct from SBS4, which is associated with lung and head and neck cancers, and SBS92, which is found in bladder cancer in the context of tobacco smoking (*27*). A high prevalence of smokeless tobacco usage has been reported in athletes (*31, 32*); smokeless tobacco usage in American football players is unknown. SBS32 is associated with azathioprine treatment and was recently reported to be strongly associated with age in oligodendrocytes with an absence in aging neurons (*23*). Other COSMIC signatures, including SBS30 and SBS89, were present in controls but increased in CTE; SBS30 is associated with deficient base excision repair which may also play a role in the excessive DNA damage.

### Somatic Indels in CTE neurons

Compared to SNVs, somatic Indels showed a more pronounced increase in CTE than in controls (*P* = 0.002, LME model; *P* = 2.8 × 10^-8^, two-tailed Wilcoxon test; Fig. 3A, B). For RHI neurons, as with somatic SNVs, we did not observe any significant difference in somatic Indel burden compared to controls, further supporting our hypothesis that increased somatic burden is specific to CTE pathology. Notably, CTE neurons accumulated a mean of 313 excess Indels per cell versus controls, approximately three times the CTE-specific SNV increase and equivalent to Indels accumulated over > 100 years (given the normal neuronal rate of 2-3 somatic Indels per year per genome). These CTE excess Indels are predominantly 2 to 4 bp deletions (Fig. 3C).

**Fig. 3.**
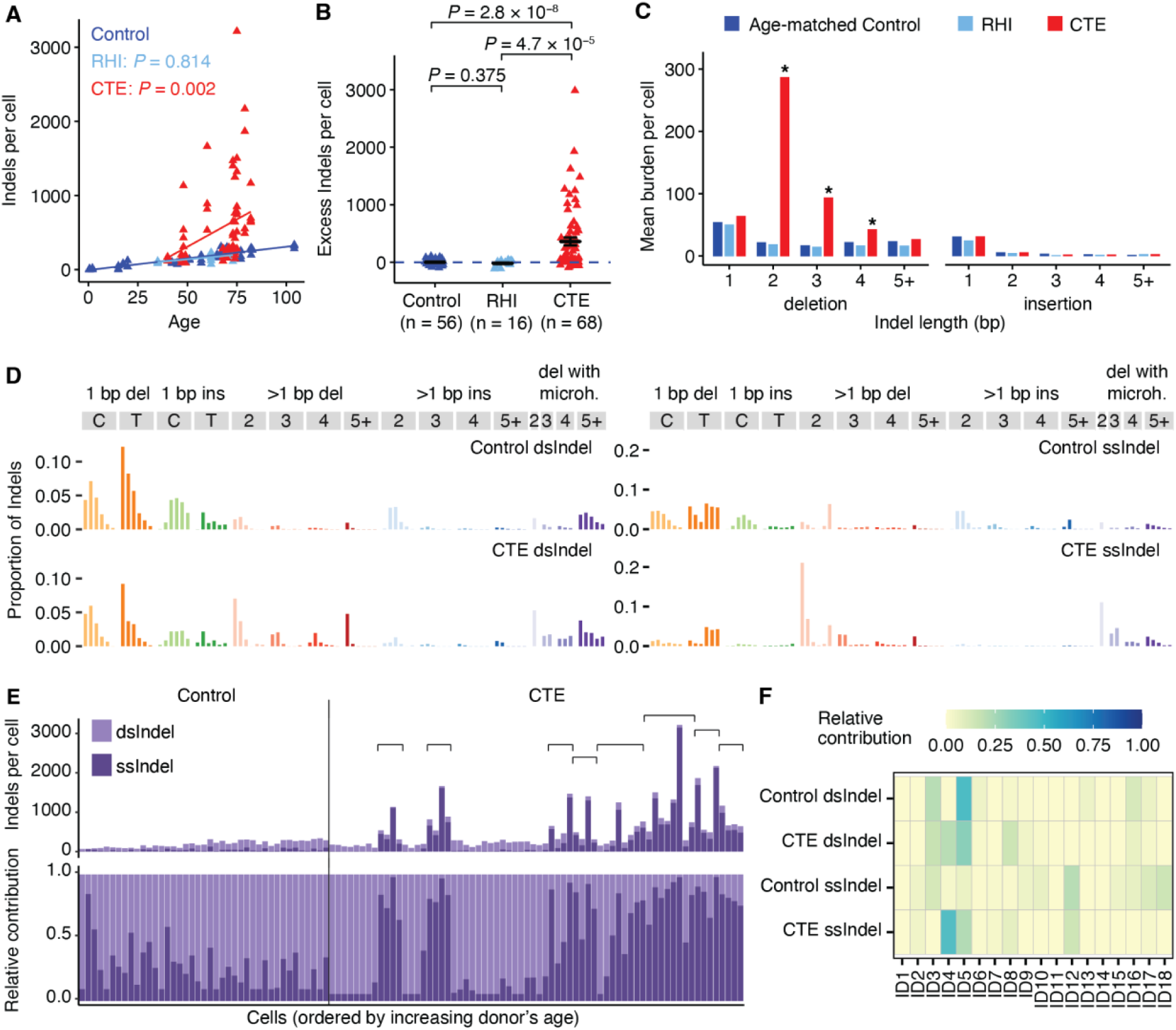
Somatic Indel burden and mutational signatures in CTE neurons. Somatic Indel burdens were identified from PTA scWGS data, with strand-related analysis inferred from mutational signatures extracted from META-CS scWGS data. (**A**) Increased somatic Indel burden in CTE (red) brains compared to RHI (light blue) brains and neurotypical controls (dark blue). Somatic Indel burden (from each neuron as a triangle) estimated by SCAN2 is fitted against age by clinical conditions using a LME model (neurotypical control: dark blue; RHI: light blue, *P* = 0.814; CTE: red, *P* = 0.002). (**B**) CTE neurons show a significant excess of somatic Indel burden (*P* = 2.8 × 10^-8^, two-tailed Wilcoxon test) while RHI neurons show no significant difference (*P* = 0.375, two-tailed Wilcoxon test) compared to neurotypical controls. Data is mean ± standard error. The dashed blue line shows Indels attributable to age (zero excess). (**C**) Comparison of all types of Indels across age-matched controls, RHI, and CTE. Data is mean burden per cell. Asterisk denotes significant changes in certain types of Indels when compared to age-matched controls (*P* < 0.05, two-tailed Wilcoxon test). 2 to 4 bp deletions are significantly more abundant in CTE neurons but not in RHI neurons. (**D**) Double-stranded (ds) and single-stranded (ss) Indel signatures in neurotypical controls and CTE extracted from META-CS data. (**E**) Absolute (top) and relative (bottom) contribution of dsIndel and ssIndel signatures in PTA-profiled neurotypical control and CTE neurons. The pair of dsIndel and ssIndel signatures used for decomposition is determined by the clinical condition of PTA neurons. We observe a pronounced ssIndel contribution in neurons from certain CTE cases (cases are indicated by brackets). (**F**) Decomposition of dsIndel and ssIndel signatures to COSMIC ID database, where we see a pronounced presence of ID4 specific to CTE.

We developed a new computational pipeline (see Methods) specifically for identifying somatic Indels in single-cell META-CS data and extracted mutational signatures that represented double-stranded and single-stranded Indels (dsIndels and ssIndels) in CTE and neurotypical controls (Fig. 3D). We confirmed the robustness of these signatures with different calling thresholds in META-CS (fig. S8A-D). To assess the contribution of double-stranded mutations versus single-stranded lesions in PTA neurons, we decomposed the Indel spectrum of each PTA neuron into a pair of dsIndel and ssIndel signatures matched with its clinical group. Interestingly, we found that the excess somatic Indels in some CTE individuals are primarily single-stranded, whereas other CTE individuals and controls have more modest numbers of Indels that are primarily double-stranded (Fig. 3E).

To explore the potential biological processes that gave rise to dsIndels and ssIndels in CTE neurons, we decomposed META-CS-derived signatures into the COSMIC reference Indel (ID) signatures (v.3.2) using NMF (Fig. 3F). Some signatures were present in both CTE and neurotypical controls. ID5, a clock-like signature associated with aging, contributes to dsIndels of both groups, as well as ssIndels of CTE. There is also substantial contribution from ID3 in dsIndels of both groups which is associated with tobacco smoking and healthy aging (*17*). ID12, a signature with unknown etiology where Indels occur primarily in repetitive regions, showed a strong contribution to ssIndels of both groups, implying that repetitive regions may be prone to single-stranded lesions or sequencing errors, which are difficult to distinguish with the current technology. ID8, mainly present in CTE, has a clock-like correlation with age both in cancers and healthy neurons (*23*). Notably, ID4, mainly characterized by 2 to 4 bp deletions, has a predominant and robust (fig. S8E-H) presence in CTE and is minimal in controls, consistent with the bias towards 2 to 4 bp deletions we observed in CTE PTA neurons (Fig. 3C). While the signal is more pronounced in the ssIndel signature, the dsIndel signature also showed a substantial ID4 contribution, which indicates a mutagenic process where a subset of ID4-associated DNA lesions become double-stranded mutations.

### Somatic ID4-like deletions in certain CTE individuals

Case-by-case inspection revealed that the excess somatic Indels in CTE were driven by a subset of individuals (Fig. 4A). Some CTE individuals consistently harbored more Indels than others (mean excess Indels per cell > 200 compared to age-matched controls), designated High-Indel CTE. In contrast, some CTE individuals exhibited much lower excess Indels compared to age-matched controls, designated Low-Indel CTE. This distinction in excess Indels between High-Indel and Low-Indel CTE groups remained after controlling for covariates representing sample and sequencing quality (fig. S9). While individuals in the High-Indel group were generally older (> 70 years old), two younger High-Indel individuals suggest that other factors beyond age may contribute. Interestingly, we found a significant association between somatic Indel burden in CTE and the duration of symptoms (i.e., the time between symptoms onset and death; *P* = 0.022, LME model), which was stronger than the association with age at death (*P* = 0.059, LME model), suggesting that the duration of disease contributes to excess somatic Indels. The *APOE* ε4 allele has been identified as a prominent risk factor for AD (*33*) and is significantly associated with the severity of CTE tau pathology (*34*), however, we found no significant association of *APOE* ε4 with somatic Indel burden in CTE after controlling for age (*P* = 0.503, 0.786, and 0.766 for dominant, additive, and recessive genetic models, LME model). Years of playing football, a proxy for cumulative RHI exposure, also did not show a significant association (*P* = 0.576, LME model). This further supports that, while the duration of RHI exposure may increase the risk and severity of CTE (*6, 24–26*), it does not significantly affect somatic mutation accumulation in CTE, consistent with our findings in SNV. We then re-analyzed PTA AD neurons from our previous study (*18*) and noted a similarly high rate of somatic Indels in certain AD individuals (designated High-Indel AD versus Low-Indel AD, Fig. 4B). This indicates that excessive somatic Indels may contribute to multiple tau-based neurodegenerative diseases.

**Fig. 4.**
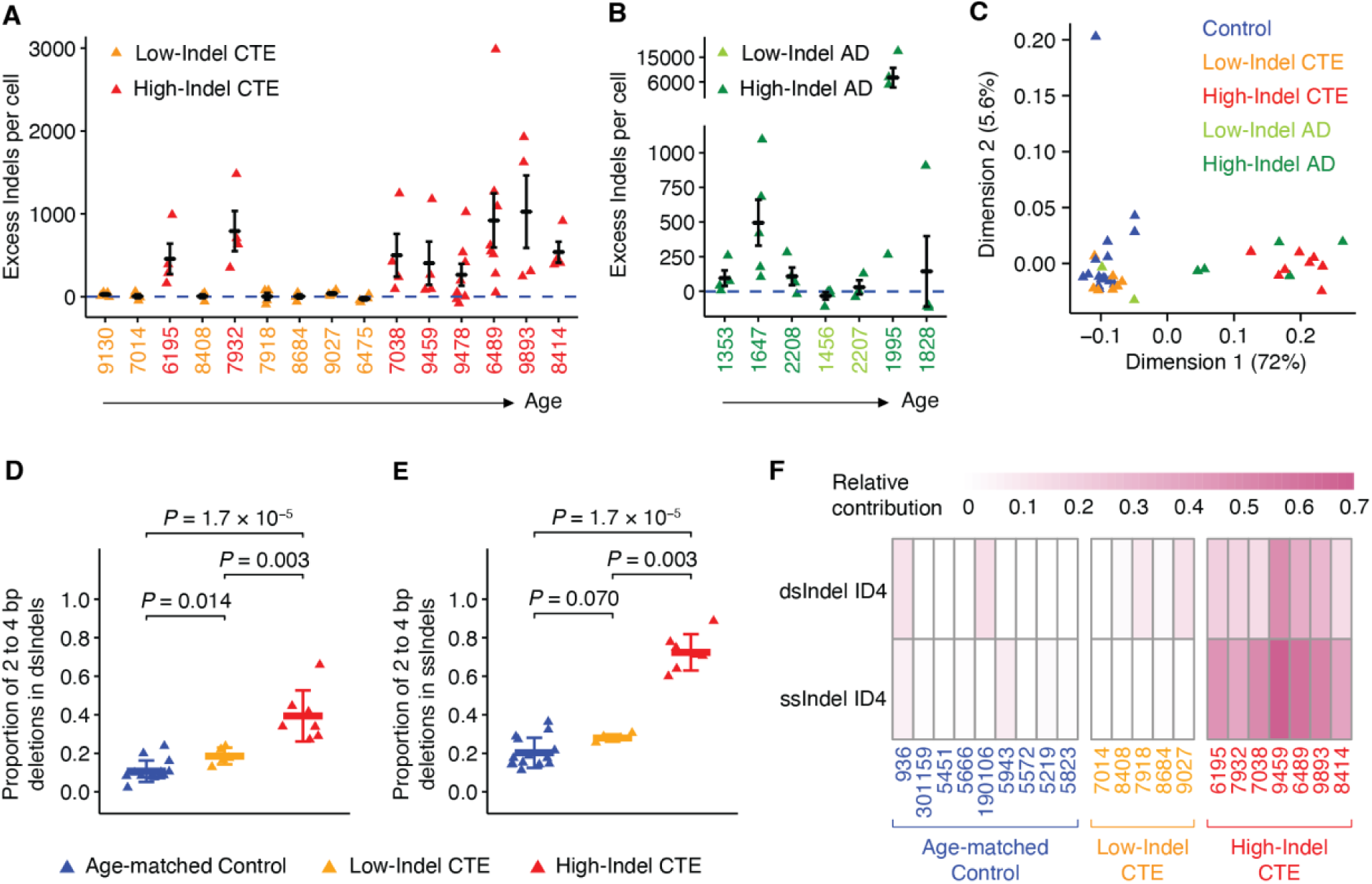
Somatic ID4-like deletions in certain CTE individuals. (**A**, **B**) Excess somatic Indels in each CTE (A) and AD (B) case ordered by increasing age. CTE and AD cases are separated into either High-Indel group (High-Indel CTE: red, High-Indel AD: dark green) or Low-Indel group (Low-Indel CTE: yellow, Low-Indel AD: light green) based on whether they have a significant excess of Indels (> 50). Data is mean ± standard error. The dashed blue line shows Indels attributable to age (zero excess). (**C**) Principal component analysis (PCA) clustering of Indels from each case across neurotypical controls, CTE, and AD. The first two dimensions are used for visualization with the percentage of variance explained shown for each dimension. Cases with < 15 Indels are filtered out. (**D**, **E**) Comparison of proportions of 2 to 4 bp deletions in dsIndels (D) and ssIndels (E) across age-matched controls, Low-Indel CTE, and High-Indel CTE. Each triangle represents an individual from META-CS data. High-Indel CTE showed a significantly higher proportion in both dsIndels and ssIndels (*P* = 1.7 × 10^-5^, two-tailed Wilcoxon test), whereas Low-Indel CTE showed marginally significant difference in dsIndels (*P* = 0.014, two-tailed Wilcoxon test) compared to age-matched controls. Data is mean ± standard deviation. (**F**) Relative contribution of ID4 to dsIndels and ssIndels of each case from META-CS data shown as a heatmap. Case IDs are colored by their group assignment (age-matched control: dark blue, Low-Indel CTE: yellow, High-Indel CTE: red).

The High-Indel CTE and AD individuals present a distinct mutational pattern. Unsupervised clustering of PTA Indel spectra identified High-Indel CTE and AD individuals as a separate cluster from Low-Indel CTE and AD individuals and neurotypical controls (Fig. 4C, fig. S10). The same clustering results were reinforced by dsIndel and ssIndel spectra from META-CS data (fig. S11). The distinct pattern in High-Indel CTE and AD individuals is mainly characterized by 2 to 4 bp deletions, where the proportion of 2 to 4 bp deletions is significantly higher in High-Indel CTE compared to age-matched controls for both dsIndels and ssIndels (*P* = 1.7 × 10^-5^, two-tailed Wilcoxon test, Fig. 4D-E). In line with the fact that ID4 comprises primarily 2 to 4 bp deletions, we observed a consistent contribution of both double- and single-stranded ID4 across all High-Indel CTE (Fig. 4F). As shown in fig. S12, High-Indel CTE demonstrated a significant increase of ID4 contribution for both dsIndels and ssIndels compared to age-matched controls (*P* = 0.001 and 1.7 × 10^-4^, two-tailed Wilcoxon test), whereas Low-Indel CTE had similar ID4 contribution as age-matched controls in dsIndels and ssIndels (*P* = 0.286 and 0.092, two-tailed Wilcoxon test). This highlights the role of ID4-associated biological processes in elevated DNA damage in CTE. The etiology of ID4 is largely undetermined; however, a recent study (*35*) demonstrated that ID4 may be mediated by topoisomerase 1 (TOP1) activity through a mechanism wherein TOP1 cleaves one of the DNA strands and leads to strand realignment to produce dinucleotide deletions. This model aligns with our observation that ID4 was more pronounced in single-stranded events. However, it is unclear how TOP1 activity may interact with CTE pathology and neurodegeneration.

### Influence of transcriptional activity and chromatin accessibility on somatic mutation in CTE

Gene transcription and epigenetic state are closely linked to DNA damage and repair in neurons (*36, 37*). Previous studies have reported that both gene expression and chromatin accessibility are positively correlated with somatic mutation burden in healthy neurons (*17, 18, 23*). To investigate whether somatic mutations have distinct enrichment patterns in CTE neurons, we generated single-nucleus RNA sequencing (snRNA-seq) data from CTE and neurotypical control samples (fig. S13) and utilized published single-nucleus assay for transposase-accessible chromatin with sequencing (snATAC-seq) data from sample-matched neurotypical controls (*23*). We extracted neuronal profiles to eliminate potential biases from other cell types.

We confirmed an increasing trend of somatic SNV burden with gene expression and chromatin accessibility in both control and CTE (Fig. 5A). Using Signatures A and C to further dissect the enrichment patterns according to mutational processes (Fig. 5B), we observed a positive correlation between gene expression and somatic SNVs attributed to Signature A for both control and CTE, while Signature C showed a negative correlation (Figure 5B), consistent with previous findings in AD neurons (*18*) and reinforcing the shared findings between CTE and AD. As for signature-specific enrichment in open chromatin, we found a positive correlation between Signature A-related SNVs and accessibility levels, but no clear trend for Signature C-related SNVs, which may be explained by complex relationship between chromatin accessibility and gene expression activity in highly expressed genes (fig. S14) as well as the potential confounding of global genome (GG) NER-related SNVs in regulatory elements, as Signature C was identified in genetic disorders with deficiency in either TC-NER or GG-NER (*21*). For somatic Indels, we separately extracted double- and single-stranded signatures from High-Indel CTE, Low-Indel CTE, and controls, and found that total burden as well as burden attributed to either signature showed a moderately positive correlation with both gene expression and chromatin accessibility (Fig. 5C, D). The similar pattern between dsIndel and ssIndel suggests that they may share a similar mutagenesis mechanism, as single-stranded lesions are an intermediate stage before either being repaired or fixed into double-stranded mutations. Moreover, the similarity across three groups suggests that different Indel patterns, particularly ID4-like patterns, are not differentially affected by transcriptional activity or chromatin accessibility.

**Fig. 5.**
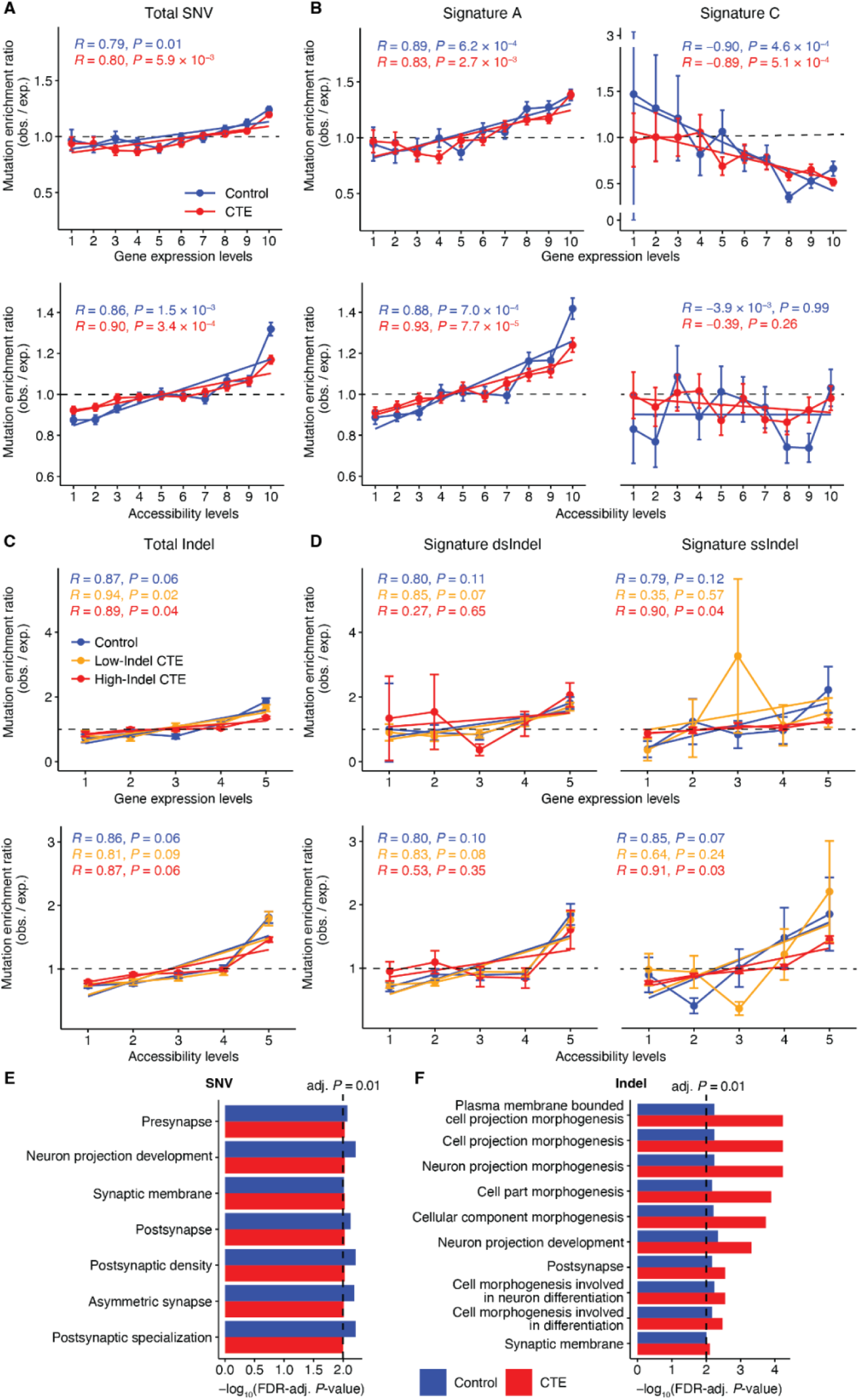
Enrichment analysis of somatic SNVs and Indels in CTE neurons. (**A**–**D**) Somatic SNV (A, B) and Indel (C, D) density as a function of gene expression and chromatin accessibility levels. Neuronal transcriptional profiles were characterized from snRNA-seq data sequenced in this study. Neuronal accessibility profiles were obtained from snATAC-seq data in Ganz *et al.*(*23*). Genes and open chromatin regions are separated into 10 (for SNV) or 5 (for Indel) equally sized groups with increasing levels indicating increasing expression or accessibility. Observed density of each expression or accessibility group is obtained by overlapping the original mutation call sets with regions of each group. Expected density is calculated from 1000 permutations of mutation call sets overlapped with regions of each group. Enrichment ratio is calculated by observed / expected density for each permutation, and mean ratio over 1000 permutations is used to construct the trend line by linear model (error bar indicates standard deviation). Pearson correlation coefficient (*R*) and two-tailed p-value (*P*) are shown. obs.: observed, exp.: expected. (**E**, **F**) Gene ontology (GO) analysis of genes where somatic SNVs (E) and Indels (F) are located. GO terms with FDR-adjusted *P* < 0.01 in both CTE and control are reported.

We further examined the distribution of somatic mutations in protein-coding genes for their potential functional impact. Gene ontology (GO) analysis of genes mutated with somatic SNVs (Fig. 5E) and Indels (Fig. 5F) revealed a consistent enrichment in neuronal functions such as synaptic structures and neuron projection. This observation aligns with our finding that somatic mutations are enriched in regions of high expression and open chromatin, reflecting the tendency of neuronal genes to be actively transcribed in neurons. Taken together with the CTE-specific somatic burden increase, our results suggest that excess somatic mutations, notably Indels in CTE neurons, may disrupt functions of essential neuronal genes and eventually contribute to neurodegeneration (Fig. 6).

**Fig. 6.**
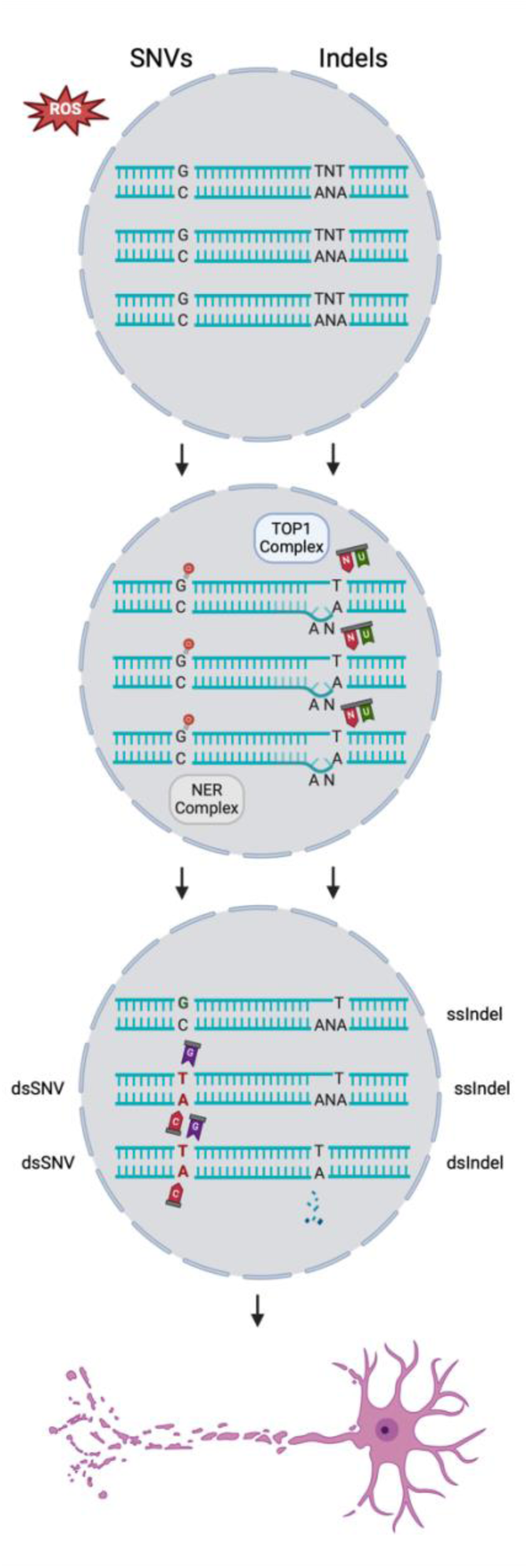
Proposed mechanisms of mutagenesis for somatic SNVs and Indels in CTE. An elevated level of DNA damage in CTE neuron in part caused by ROS leads to the accumulation of both somatic SNVs and Indels. For SNVs, oxidative damage frequently results in 8-oxoguanine on one strand which becomes a double-stranded C>A mutation when NER and other pathways fail to repair it. For Indels, genome-embedded ribonucleotides are removed by TOP1-mediated activities during which a cleavage on one strand with the TNT motif leads to strand realignment and consequently a 2 bp single-stranded deletion. A small portion of the single-stranded deletions may become double-stranded deletions potentially through other endogenous processes. The accumulation of such DNA damage may eventually lead to neurodegeneration and cell death.

## Discussion

In this study, we characterized somatic mutations in CTE neurons using two different scWGS methods and revealed significantly elevated DNA damage leading to somatic SNVs and Indels in CTE, in patterns distinct from normal aging. We propose distinct mutagenic pathways for the accumulation of somatic SNVs and Indels (Fig. 6). Excess somatic SNVs in CTE were largely contributed by Signature C, a pattern that is rare in controls and we found at a higher relative proportion in CTE compared to AD. Given the shared feature of tau pathology, CTE and AD may share common pathways for accumulating DNA damage, possibly related to oxidative damage (*18*). We also found a subset of High-Indel CTE individuals that harbored excess double- and single-stranded Indels, with a predominant contribution from ID4 (*27*). Analysis of somatic Indels from AD showed a subset with similar excessive Indels.

Previous studies have highlighted RHI as the likely cause of CTE (*38, 39*); however, not all individuals exposed to RHI develop CTE. We found that the somatic mutation burden of the RHI group was similar to neurotypical controls, implying that the significantly elevated DNA damage in CTE is distinct from long-term exposure to RHI. A limitation of this study is that the RHI group is small, and a larger group of varying RHI exposures is needed to validate this finding. Our cohort focused on severe CTE (stage III and IV); future studies that include early-stage CTE are needed to assess the role of somatic mutations in disease progression. Moreover, other unique features of CTE such as the perivascular localization of tau deposition at the depths of the cortical sulci may unveil novel linkage between DNA damage and pathogenesis. Our analysis between the sulcus and gyrus showed no significant difference (fig. S3) but statistical power was limited. A larger such study is necessary to detect more subtle differences between topological areas of the brain. In addition, we did not detect any somatic mutations in genes known to be associated with neurodegenerative tauopathies (*APP*, *PSEN1*, *PSEN2*, *APOE*, and *MAPT*), similar to a previous report in AD (*18*).

The incorporation of duplex sequencing data provided a unique opportunity to distinguish single-stranded from double-stranded genomic alterations. Surprisingly, somatic SNVs and Indels in CTE neurons appeared to have distinct distributions of double- and single-stranded events that may be individual-specific. While most somatic SNVs from all CTE individuals were represented on both strands, High-Indel CTE individuals present a distinct Indel mutational pattern contributed by many features of ID4. ID4 was recently reported to result from transcription-associated mutagenesis (*35*). The authors proposed that these events occur when ribonucleotides are mis-incorporated into DNA and subsequently cleaved by TOP1 instead of RNase H2. The TOP1 repair pathway creates short deletions with a pattern that is most similar to ID4. These double-stranded deletions are thought to arise via an intermediate step of single-stranded > 1 bp deletions, which aligns with our observation that ID4-like patterns are present in both dsIndels and ssIndels of High-Indel CTE but with a stronger signal in ssIndels. While the association between RNase H2 deficiency and neurodegenerative disease is unknown, our results raise the possibility that a large quantity of genome-embedded ribonucleotides in CTE and AD neurons might overwhelm the ribonucleotide excision repair pathway, with the TOP1-mediated process leaving large numbers of > 1 bp deletions on single DNA strands of which only a small proportion could be efficiently repaired or fixed into double-stranded Indels in post-mitotic neurons. Further investigation is required to identify the determining factors on the occurrence of these events, and why they tend to be quite abundant in some CTE and AD individuals and less common in others. It is also worth noting that ID4 may not fully explain the DNA damage in CTE, as some excess somatic Indels in CTE, particularly 2 bp deletions in one repeat unit, are limited in the ID4 spectrum. This suggests other mutagenic processes may play a role in congruence with or independent of ID4-associated processes. Nonetheless, our analysis highlights that CTE can be viewed as a genomic process in affected neurons, with progressive transcription-related accumulation of SNV and especially Indels – which accumulate to numbers equivalent to > 100 years of excess aging – creating the potential for severe dysregulation of the neuronal transcriptome (Fig. 6).

## Acknowledgments

We thank R. Mathieu and T. Berisha from the Boston Children’s Hospital Flow Cytometry Core, R. Krishnan from the Brigham and Women’s Hospital Center for Neurologic Diseases Flow Cytometry Core Facility, and R. S. Hill and J. E. Neil for their administrative help. We thank the donors of postmortem tissues for their invaluable contributions to the advancement of science.

## Funding

R56 AG079857 (A.Y.H., C.A.W., E.A.L.)

R01 AG088082 (A.Y.H.)

Alzheimer’s Association Research Fellowship (A.Y.H.)

K08 AG065502 (M.B.M.)

DP2 AG086138 (M.B.M.)

R01 AG082346 (M.B.M.)

Alzheimer’s Disease Research program of the BrightFocus Foundation A20201292F (M.B.M.)

Doris Duke Charitable Foundation Clinical Scientist Development Award 2021183 (M.B.M.)

Suh Kyungbae Foundation (E.A.L.)

DP2 AG072437 (E.A.L.)

R01 NS032457-20S1 (C.A.W.)

R01 AG070921 and AG078929 (C.A.W., E.A.L.)

Templeton Foundation (C.A.W.)

Allen Discovery Center program, a Paul G. Allen Frontiers Group advised program of the Paul G. Allen Family Foundation (C.A.W., E.A.L.)

Howard Hughes Medical Institute (C.A.W.)

National Institute of Neurological Disorders and Stroke U01NS086659, U01NS093334, U54NS115266, R01NS078337, R56NS078337, K23NS102399 (A.C.M.)

National Institute of Aging P30AG13846 (A.C.M.)

Department of Veterans Affairs I01 CX001135 (A.C.M.)

National Operating Committee on Standards for Athletic Equipment (A.C.M.)

Nick and Lynn Buoniconti Foundation (A.C.M.)

Concussion Legacy Foundation (A.C.M.)

Andlinger Foundation (A.C.M.)

WWE (A.C.M.)

NFL (A.C.M.)

## Author contributions

M.B.M., A.C.M., and C.A.W. conceived the study, further developed by C.C.M. C.C.M. and M.B.M. performed tissue dissection, single-nuclei sorting, PTA genome amplification, and oversaw sequencing with assistance from S.M.N., S.L.K., and G.A.M. S.M.N. performed META-CS genome amplification, with assistance from C.C.M. and M.B.M. G.D. performed data analysis with assistance from S.M. and J.K. G.D. developed and applied the software for mutation calling in META-CS data. A.C.M., M.U., J.D.C., and E.S. provided brain samples, clinico-pathological data, and assisted with data interpretation. G.D., C.C.M., S.M., and K.S.B. prepared display items. G.D. and C.C.M. wrote the initial draft of the manuscript with assistance from M.B.M., A.Y.H., S.M., and K.S.B. M.B.M., A.Y.H., and E.A.L. oversaw experiments and analyses. C.A.W., E.A.L., M.B.M., and A.Y.H. supervised the study and edited the manuscript.

## Data and materials availability

Sequencing data generated in this study will be deposited in a public repository, with controlled use conditions set by human privacy regulations. All the code will be made available on Github. Other materials are available from the authors upon reasonable request.

## Supplementary Materials

Materials and Methods

Figs. S1 to S14

Tables S1 to S6

References (40–54)

## Materials and Methods

### Data reporting

No statistical methods were used to predetermine sample size. Experimenters were not blinded and experiments were not randomized.

### Human tissue samples and selection of cases of CTE

Post-mortem frozen human tissues were obtained from the UNITE Brain Bank (formerly VA-BU-CLF) at Boston University, the Massachusetts Alzheimer’s Disease Research Center (MADRC) at Massachusetts General Hospital and the NIH Neurobiobank at the University of Maryland Brain and Tissue Bank (UMBTB).

Tissue collection and distribution for research and publication was conducted according to protocols approved by the Boston University-Veterans Affairs Institutional Review Board (for BU-VA: S07-02-0087), MassGeneral Brigham Institutional Review Boards (for MADRC: 1999P009556/MGH, expedited waiver category 5) and the University of Maryland Institutional Review Board (for UMBTB: 00042077), and after provision of written authorization and informed consent. Research on these de-identified specimens and data was performed at Boston Children’s Hospital with approval from the Committee on Clinical Investigation (S07-02-0087 with waiver of authorization, exempt category 4) and at Brigham and Women’s Hospital with approval from the MassGeneral Brigham Institutional Review Board (2019P003790 for secondary use as non-human subjects research).

Neurotypical control cases had no clinical history of dementia or other neurological disease, and many were obtained as part of a previous study (*17, 18*). CTE cases were pathologically confirmed stage III-IV (*3, 40*) with no other notable neurodegenerative pathology. AD cases had a clinical history of dementia consistent with AD, pathologically confirmed AD pathological change (Braak stage V–VI) and no other notable neurodegenerative pathology.

### Isolation of individual pyramidal neurons for single-cell studies through FANS

Single neuronal nuclei were isolated using a sucrose cushion and fluorescence-activated nuclear sorting (FANS) for the neuronal nuclear transcription factor (NeuN), as described in previous work (*18*).

In brief, nuclei were prepared from unfixed frozen (at −80 °C) human brain tissue in a dounce homogenizer using a chilled tissue lysis buffer (10 mM Tris-HCl, 0.32 M sucrose, 3 mM Mg(OAc)_2_, 5 mM CaCl_2_, 0.1 mM EDTA, 1 mM DTT, 0.1% Triton X-100, pH 8) on ice. Lysed tissue was then layered on top of a sucrose cushion buffer (1.8 M sucrose 3 mM Mg(OAc)_2_, 10 mM Tris-HCl, 1 mM DTT, pH 8) and ultra-centrifuged for 1 hour at 30,000 × g. Nuclear pellets were resuspended in ice-cold PBS supplemented with 3 mM MgCl_2_, filtered, then stained with DAPI and anti-NeuN (RBFOX3) antibody directly conjugated to Alexa Fluor 488 (AF488) (Millipore MAB377X, clone A60, 1:1,250). DAPI staining allowed for isolation of single diploid neuronal nuclei apart from debris and doublet doplets. Previous work showed that NeuN staining produced a bimodal signal distribution (*18*). Large neuronal nuclei, representing excitatory pyramidal neurons, were then identified by flow cytometry (using software BD FACSDiva v.8.0.2) by targeting the nuclei with highest NeuN signal among the NeuN+ neuronal fraction, while also gating for the population with the highest forward scatter area (FSC-A) signal (*18*). As we reported previously, this high-FSC-A, high-NeuN population represents large pyramidal neurons, which are 99.3% excitatory neurons (*18*), comprising 2–5% of the total population of nuclei in each sample.

### snRNA-seq of CTE and control large neurons

Single-nucleus RNA sequencing (snRNA-seq) was used to assess the mutational enrichment and composition of a population of large neurons in CTE along with control cases. snRNA-seq of these two populations of cellular nuclei was performed on representative tissue samples (control individual UMB1465, prefrontal cortex; CTE individual CTE9310, prefrontal cortex). Nuclei were isolated as described above, with the following modifications: 0.2 U μl−1 Protector RNAse inhibitor (Roche RNAINH-RO) and 0.2 U μl −1 SuPERase-IN RNAse inhibitor (Invitrogen) were both added to the tissue lysis buffer and to the immunostaining buffer, and MgCl2 was omitted [check] from the immunostaining buffer. For each of these populations, ∼16,000 nuclei were sorted into one well of a 96-well plate, then subjected to snRNA-seq using the 10X Genomics Next GEM Single Cell 3′ GEM Kit v3.1 and Chromium Controller. From these two populations, two libraries were prepared, each with dual indexes using the 10X Genomics Dual Index Plate. Each library was then sequenced on Illumina.

### scWGS of pyramidal neurons using PTA

Single neurons, prepared as described above, were sorted one nucleus per well into 96-well plates and their genomes were amplified by PTA (*22*), a method that pairs an isothermal DNA polymerase with a termination base to induce quasi-linear amplification. PTA reactions were performed using the ResolveDNA Whole Genome Amplification Kit (BioSkryb Genomics). Nuclei were sorted into 3 μl Cell Buffer pre-chilled on ice. Nuclei were then lysed by addition of 3 μl MS Mix, with mixing at 1,400 rpm performed after each step. Lysed nuclei were then neutralized with 3 μl SN1 buffer, followed by 3 μl of SDX reagent and a 10-min. incubation at room temperature. Next, 8 μl of reaction mix (containing polymerase) was then added, for a total reaction volume of 20 μl. Amplification was carried out for 10 h at 30 °C, followed by enzyme inactivation at 65 °C for 3 min. Amplified DNA was then cleaned up using AMPure, and the yield was determined using PicoGreen binding (Quant-iT dsDNA Assay Kit, Thermo Fisher Scientific). Samples were then subjected to quality control by multiplex PCR for four random genomic loci as previously described (*21*). Amplified genomes showing positive amplification for all four multiplex PCR loci were prepared for Illumina sequencing.

Libraries were prepared following a modified KAPA HyperPlus Library Preparation protocol described in the ResolveDNA EA Whole Genome Amplification protocol. In brief, end repair and A-tailing were performed for 100-500 ng amplified DNA input. Adapter ligation was then performed using the SeqCap Adapter Kit (Roche, 07141548001). Ligated DNA was cleaned up using AMPure and amplified through an on-bead PCR amplification. Amplified libraries were selected for a size of 300–600 bp using AMPure. Libraries were subjected to quality control using PicoGreen and TapeStation HS DS100 Screen Tape (Agilent PN 5067-5584) before sequencing. Single-cell genome libraries were sequenced on the Illumina NovaSeq platform (150 bp × 2) at 30× coverage (table S1). Data from PTA-amplified neuronal genomes in CTE were analyzed alongside previously reported data from control (*17*) and AD neurons (*18*).

### scWGS of pyramidal neurons using modified META-CS

The genomes of single neuronal nuclei were amplified by META-CS [19], a transposase-based whole genome amplification technique in which each DNA fragment is tagged and barcoded with 16 unique tags (Dataset S1 of Xing *et al*. (*19*)), allowing for single-cell, strand-resolved identification.

DNA oligos were ordered from IDT. Each of the 16 META-CS transposons were annealed and assembled into transposomes with Diagenode Tagmentase (Diagenode; C01070010) per manufacturer’s protocol and stored at −80 °C.

Single neuronal nuclei, isolated as previously described, were sorted one per well into 96 well plates, and lysed in 2 μl of 1x Single Cell Lysis Buffer (20 mM Tris, pH 8.0, 20 mM NaCl, 0.15% Triton X-100, 25 mM dithiothreitol, 1 mM EDTA, 1.5 mg/mL Thermolabile Proteinase K (TLPK) (NEB, P8111S)) at 30 °C for 1 h, 55 °C for 10 min. Single cell lysates were stored at −20 °C if not immediately amplified.

Lysed nuclei were then transposed by the addition of 8 μL transposition mix (5ul Diagenode 2X Tagmentation buffer (Diagenode; C01019043), 1 μl diluted META-CS transposome, 2 μl H_2_O), mixed at 1640 rpm for 1 min, spun down at 1500 rpm, and incubated at 55 °C for 15 min. Transposases were removed by the addition of 2 μL 6X stop buffer containing 300 mM NaCl, 45 mM EDTA, 0.01% Triton X-100, and 1 mg/mL TLPK, with mixing and incubation at 37 °C for 30 min, 55 °C for 10 min.

First-strand tagging was performed by the addition of 13 μL Strand Tagging Mix 1 containing 5 μL Q5 reaction buffer (NEB; B9027S), 5 μL Q5 high GC enhancer (NEB; B9028A), 0.85 μL 100 μM (total) Adp1 primer mix (Dataset S1 of Xing *et al*. (*19*)), 0.6 μL 100 mM MgCl2, 0.55 μL water, 0.5 μL 10 mM (each) dNTP mix (Thermo Scientific; R0192), 0.25 μL of 20 mg/mL bovine serum albumin (NEB; B9000S), and 0.25 μL Q5 DNA polymerase (NEB; M0491S), followed with mixing and incubation at 72 °C for 3 min, 98 °C for 30 s, 62 °C for 5 min, 72 °C for 1 min. ADP1 primers were removed with 1 μL Thermolabile Exonuclease I (NEB; M0568L), with mixing and incubation at 37 °C for 15 min, 65 °C for 5 min.

Second-strand tagging was performed by the addition of 4 μL Strand Tagging Mix 2 containing 1 μL Q5 reaction buffer, 1 μL Q5 high GC enhancer, 0.95 μL 100 μM (total) Adp2 primer mix (Dataset S1 of Xing *et al*. (*19*)), 0.85 μL water, 0.1 μL 10 mM (each) dNTP mix, and 0.1 μL Q5 DNA polymerase, and incubation at 72 °C for 3 min, 98 °C for 30 s, 62 °C for 5 min, 72 °C for 1 min. Adp2 primers were removed by repeating the exonuclease step described above.

Strand tagging products were amplified by the addition of 19 μL PCR mix containing 1 μL NEBNext Multiplex Oligos Universal Primer, 1 μL NEB Index Primers (NEB; E7335S, E7500S, E7710S, E7730S), 4 μL Q5 reaction buffer, 4 μL Q5 high GC enhancer, 0.4 μL 10 mM (each) dNTP mix, 8.4 μL water, and 0.2 μL Q5 DNA polymerase and incubation at 98 °C for 20 s, 13 cycles of [98 °C for 10 s, 72 °C for 2 min], 72 °C for 2 min.

Single-cell libraries were pooled together, and then libraries were purified by DNA Clean and Concentrator-5 columns (Zymo; D4013) and amplification efficiency was checked for fragment size and concentration by Agilent TapeStation or Agilent Bioanalyzer. Size selection was performed with Ampure XP beads (Beckman Coulter; A63880), wherein the pooled library was divided into three groups based on fragment size. Medium-size fragments (∼300 - 1000 bp) were selected first by the addition of 0.6X beads then by a further addition of 0.25X beads (for a final 0.75X).

### Read mapping and BAM file generation for bulk and PTA data

BWA (v0.7.15) (*41*) was first used to map reads from bulk WGS and PTA scWGS data onto the human reference genome (GRCh37 with decoy) with default parameters. Then, duplicate reads were marked by MarkDuplicates of Picard (v.2.8.0), followed by local Indel realignment and base quality score recalibration using Genome Analysis Toolkit (GATK) (v.3.5) (*42*) to generate BAM files for mutation calling.

### Quality measures of single-cell genome amplification

To evaluate the quality of single-cell genome amplification for both PTA and META-CS data, we used a number of measures. First, we calculated the median absolute pairwise differences (MAPD) to quantify the evenness of amplification as described previously (*43*). MAPD score was computed by binning the genome, estimating copy number of each bin, and taking the median of absolute pairwise differences between log2-transformed copy number ratios of adjacent bins. A lower MAPD score indicates more even amplification of the genome. Further, to account for the variance of the copy number ratio distribution, we calculated the coefficient of variation (CoV) of these absolute pairwise differences by taking the ratio of their standard deviation to their mean. Sequencing depth was estimated using the total number of properly mapped and paired reads (from samtools stats) multiplied by read length and divided by the whole genome length. To account for amplification bias, we estimated the allelic and locus dropout rates using a set of high-quality germline heterozygous SNPs that overlap with common variants from the 1000 Genome Project. Allelic dropout sites have either REF depth or ALT depth < 2. Locus dropout sites have total depth < 5. Strand dropout rate was estimated as the square root of allelic dropout rate.

### Calling of somatic SNVs and Indels from PTA data

We used Single Cell ANalysis 2 (SCAN2, v.1.0) (*17*) to identify somatic SNVs and Indels from single-cell PTA data with matched bulk data. First, we generated four cross-sample panels. One panel for 17 control individuals from UMBTB, one panel for 4 RHI individuals, one panel for 15 CTE individuals and 2 control individuals from UNITE Brain Bank (formerly VA-BU-CLF), and one panel for 7 AD individuals. Each panel was configurated by running “scan2 config −-analysis makepanel” with following reference parameters, human reference genome GRCh37 with decoy (--ref), dbSNP v138 common variants (--dbsnp), and 1000 Genomes Phase 3 SHAPEIT2 phasing panel (--shapeit-refpanel). Each panel was then built by running “scan2 makepanel”. After panel generation, mutation calling was performed for each individual by running “scan2 config −-analysis call_mutations” with the same reference parameters above, each PTA BAM (--sc-bam), matched bulk BAM (--bulk-bam), and the corresponding cross-sample panel (--cross-sample-panel), followed by “scan2 run”. For each single cell, SCAN2 generated both mutation calls and genome-wide somatic mutation burden estimations (autosomes only) which were used for the subsequent PTA analyses described below.

### Preprocessing and filtering of META-CS data

Our pipeline to preprocess META-CS data and perform somatic SNV calling shares a core workflow with the previously reported method (*19*) and includes extra steps to further strengthen accuracy and remove false positives which better accommodates our modified experimental protocol. First, paired-end reads were preprocessed by pre-meta including identifying transposon barcodes, merging overlapping read ends, and trimming Illumina adapters. Then, two aligners, BWA-MEM (v.0.7.17) (*44*) and Minimap2 (v.2.12) (*45*), were used to map reads to the human reference genome (GRCh37 with decoy) and generate BAM files. Of note, since each original DNA fragment was tagged by a pair of transposon barcodes, we split BAM files by barcode pairs before running mutation calling in each barcode pair BAM. This allows us to filter out reads without matching barcodes and ensure that calling was performed on a single-molecule level (i.e. for each allele). In addition, given the relatively small number of unique barcodes, there is a chance of barcode collision where different DNA fragments are tagged by the same barcode pair. Therefore, we extracted Tn5 cut sites from each read pair, with the assumption that reads amplified from the same DNA fragment should share the same Tn5 cut sites. Other quality metrics were obtained in the same way as PTA data (described above). Due to batch effects and imbalanced coverage among pooled cells, cells with an average insert size < 250 or > 500, or a standard deviation of insert size > 750 were filtered out.

### Calling of somatic SNVs from META-CS data

We generated somatic SNV candidates by identifying variants that have no non-reference (ALT) allele read from bulk but at least four total ALT reads and at least two ALT reads from each strand for duplex support in the cell. To achieve a robust and accurate calling, we filtered out candidate sites that satisfied any of the following criteria: overlapping with the low-quality regions as previously reported (*19*), overlapping with gnomAD SNPs of ≥ 1% population frequency, having the ALT allele balance below 20%, within 100 bp from another candidate site, having reads with different Tn5 cut sites. Passing variants need to have at least 4 ALT reads in total with at least 2 ALT reads from either strand (a4s2) as well as a VAF > 0.2.

### Calling of somatic Indels from META-CS data

We established a new pipeline for somatic Indel calling from META-CS data by introducing novel modules to expand on the somatic SNV pipeline (manuscript in preparation). First, as the majority of false positive calls originated from incorrectly merged read pairs during preprocessing, we created a module to tag the merged reads and calculate a genomic window where the merging occurred. Second, we generated an additional BAM file without read merging and filtered out any candidate sites that either overlapped with the read merging window or were not present in the BAM without read merging. Other filtering criteria were the same as the ones used for somatic SNV calling, except that we used gnomAD Indel sites of ≥ 1% population frequency. Passing variants need to have at least 4 ALT reads in total with at least 2 ALT reads from either strand (a4s2) as well as a VAF > 0.2.

### Calling of single-stranded variants from META-CS data

Single-stranded SNV and Indel calls were detected in a similar fashion as double-stranded calls described above. We generated candidate sites by identifying variants that have no ALT read from bulk, at least four ALT reads from the variant strand in the cell, and no ALT read from the non-variant strand in the cell. Then, the same filtering strategy except for the ALT allele balance was implemented to generate the final call set. Passing variants need to have at least 4 ALT reads from the variant strand and at least 4 REF reads from the non-variant strand with VAF > 0.2.

### Additional filtering of SNV and Indel calls from META-CS

To account for batch effects and technical artifacts, we implemented a set of filters to remove false positives. The same filters were applied to both double- and single-stranded SNVs and Indels. We removed variant sites that meet at least one of the following criteria: 1) variant site is covered by more than one barcode-pair, 2) barcode-pair covering the variant site has > 1 set of Tn5 cut sites, 3) variant site located within 20 bp of the read ends.

### Linear mixed-effects model of somatic mutation burden

We used linear mixed-effects (LME) models from the lme4 (v.1.1-27.1) R package (*46*) to investigate potential associations between somatic mutation burden and other covariates of interest. Somatic mutation burden was modeled as a continuous outcome, covariates of interest including age, disease status, and cell quality measures were modeled as fixed effects, and individuals were modeled as random effects due to potential correlations between cells from the same individual. P-values of fixed effects were obtained from t-tests with the Satterthwaite approximation using the lmerTest (v.3.1-3) R package (*47*). To calculate QC-corrected burden, we first modeled each quality measure and age as fixed effects in control neurons to obtain the coefficient of this quality measure’s contribution, and then subtracted its contribution from total somatic burden of each cell before modeling the corrected somatic burden against age and disease status to generate QC-corrected burden comparisons.

### Association analysis of somatic mutation burden

We used the same LME model described above to evaluate the association between somatic mutation burden and covariates of interest in CTE individuals. For years of playing football, somatic SNV or Indel burden was modeled as a continuous outcome, age and the number of years playing football were modeled as fixed effects, and individuals were modeled as random effects. P-values of fixed effects were obtained from t-tests with the Satterthwaite approximation using the lmerTest (v.3.1-3) R package (*47*). For *APOE* ε4, we first genotyped all CTE individuals at two SNPs rs429358 and rs7412 using their bulk WGS data (6 possible genotypes are ε2/ε2, ε2/ε3, ε2/ε4, ε3/ε3, ε3/ε4, and ε4/ε4). Then, we used three genetic models of ε4 allele, dominant, additive (with linear penetrance), and recessive, to test the association between *APOE* ε4 genotype and somatic burden. Somatic burden was modeled as a continuous outcome, age and *APOE* ε4 genotype were modeled as fixed effects, and individuals were modeled as random effects. P-values of fixed effects were obtained from t-tests with the Satterthwaite approximation using the lmerTest (v.3.1-3) R package (*47*). In addition, we used one-tailed Wilcoxon test for football playing years and Fisher’s exact test to test the association between High-Indel CTE and Low-Indel CTE groups.

### Double- and single-stranded signature analysis

Given the four call sets (dsSNVs, ssSNVs, dsIndels, and ssIndels) from META-CS data, we used the NMF-based method from MutationalPatterns (v.3.0.1) R package (*30*) to extract a signature from each of the call set. To better interpret these signatures, we decomposed them using the COSMIC database (https://cancer.sanger.ac.uk/signatures) with matched mutation type.

Since these signatures were derived independently, the double- and single-stranded signatures for each mutation type were not orthogonal from each other and thus could not be used directly for refitting. Therefore, two intermediate signatures were introduced to serve as a proxy for refitting. For each mutation type, the double- and single-stranded signatures, DS and SS, were combined and normalized, from which two orthogonal intermediate signatures, IM_1_ and IM_2_, were extracted using NMF. DS and SS were then refitted to IM_1_ and IM_2_ so that they could be reconstructed as

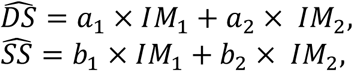

where a_1_, a_2_, b_1_, and b_2_ were proportional contributions normalized to IM_1_ and IM_2_ (i.e. a_1_ + b_1_ = 1 and a_2_ + b_2_ = 1).

To estimate the proportions of double- and single-stranded calls within the PTA call sets, for each mutation type in each cell, PTA calls were fit to the intermediate signatures,

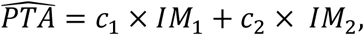

and the signature contributions were converted to describe DS and SS,

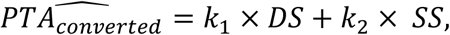

where *k*_1_ = *c*_1_ × *a*_1_ + *c*_2_ × *a*_2_ and *k*_2_ = *c*_1_ × *b*_1_ + *c*_2_ × *b*_2_. Note that all signatures and spectra are vectors.

### Known and de novo mutational signature analysis

To study the contributions of known and de novo mutational signatures to the PTA call sets, we first categorized the mutation calls from each cell into context groups predefined by COSMIC v3.2 (96 contexts for SNVs and 83 contexts for Indels). For somatic SNVs, we used MutationalPatterns (*30*) to fit the PTA calls to known signatures A and C reported in Lodato et al. (*21*) to get the proportional contribution of each signature to each cell. Then, signature-specific somatic mutation burden was calculated by multiplying the contribution by the genome-wide burden, which is comparable between cells. De novo signatures were extracted using the NMF-based method in MutationalPatterns. The number of signatures to extract was determined with 200 NMF runs based on the Kullback-Leibler divergence (method = “brunet”). PTA calls were fit to the de novo signatures in the same way as for known signatures. We used the COSMIC v3.2 database to decompose both known and de novo signatures for further interpretation.

### Permutation of somatic SNV and Indel calls

Permutation sets are crucial for robust enrichment analyses by controlling for potential biases that exist in the original mutation calls and providing powered statistical tests to evaluate the significance of enrichment. For each permutation, the original calls were randomly shuffled across the callable regions of each cell while preserving their chromosome and trinucleotide context. We generated 1000 permutation sets for each mutation type in each cell using SCAN2 by running “scan2 config −-analysis permtool” with original mutation calls (--permtool-muts), the human reference genome GRCh37 with decoy (--permtool-bedtools-genome-file), and 1000 permutations (--permtool-n-permutations), followed by “scan2 permtool”.

### Droplet-based snRNA-seq analysis

Gene count matrices for both control and CTE were acquired by aligning reads to the GRCh37 genome (v.3.0.0) using Cellranger (v.6.1.0) (*48*) with default parameters. Then we used the standard workflow from Seurat (v.4.0.5) (*49*) to process the snRNA-seq data. For the purpose of quality control, we removed genes that were not expressed in < 3 cells and cells with 1) < 200 genes, 2) < 500 counts or > 15000 counts, and 3) mitochondrial gene percentage > 5%. After applying these filters, we obtained 4619 cells for the control dataset and 4080 cells for the CTE dataset. Next, the data were normalized using the function “LogNormalize” with “scale.factor” = 10000 and scaled to a maximum value of 10. We then performed dimension reduction (principal component analysis, t-SNE, and UMAP) and cell clustering using Louvain (*50*). To identify the cell types that were captured in the snRNA-seq data, we selected marker genes for each cluster by statistical tests and assigned cell-type labels according to previously reported marker lists (*51*). Through visual evaluation, cells with the same identity were clustered together.

### Enrichment analysis with snRNA-seq and snATAC-seq data

We performed enrichment analysis of somatic mutations in transcribed gene or open chromatin regions using our in-house snRNA-seq data and previously reported snATAC-seq data generated from matching control samples (*23*). We extracted the gene expression and chromatin accessibility profiles of neurons annotated in snRNA-seq and snATAC-seq data to match the PTA data.

For gene expression enrichment analysis, we used the expression profile of control for mutation calls from control and RHI cells, and the expression profile of CTE for mutation calls from CTE cells. First, genes were ranked based on their expression and divided into equal-sized groups (10 groups for SNVs, 5 groups for Indels). Then, we intersected both original and permuted mutation calls with gene regions of each expression group. Gene regions were annotated by ANNOVAR (*52*) using the database GRCh37 refGene (*53*). Mutation calls that overlapped with multiple genes were removed. A ratio of observed to expected number of calls (i.e., enrichment ratio) was calculated for each permutation round where the observed number came from the original call set and the expected number came from the permuted call set. Enrichment was reported as the mean and standard deviation of the 1000 ratios.

For chromatin accessibility enrichment analysis, we used the processed BED file of excitatory neurons (*23*) where the genome was first separated into non-overlapping 1000 bp bins and snATAC-seq fragments were mapped to the bins. Similar to gene expression enrichment, the bins were ranked based on their coverage and divided into equal-sized groups (10 groups for SNVs, 5 groups for Indels). Then, we intersected both original and permuted mutation calls with genomic regions of each accessibility group. Enrichment ratios were calculated in the same way as above.

In addition to total somatic mutation enrichment, we performed signature-specific enrichment using the same NMF-based method for signature analysis above. For both original and permuted calls that overlapped with each expression or accessibility group, we fit the somatic SNV calls to signatures A and C and the somatic Indel calls to the double- and single-stranded signatures from META-CS. The observed/expected ratios were calculated as above.

### Gene Ontology analysis

Gene Ontology enrichment analysis was performed on genes that overlapped with somatic mutations using GOseq (v.1.42.0) (*54*) after controlling for gene length bias. A null distribution was generated by Wallenius approximation, and GO categories with less than 10 hits or more than 1000 genes were filtered out. P-values of over- and under-represented GO categories were adjusted for multiple testing using FDR. All GO categories with adjusted P < 0.05 in both CTE and neurotypical controls were reported in table S6.

---

**Fig. S1.**
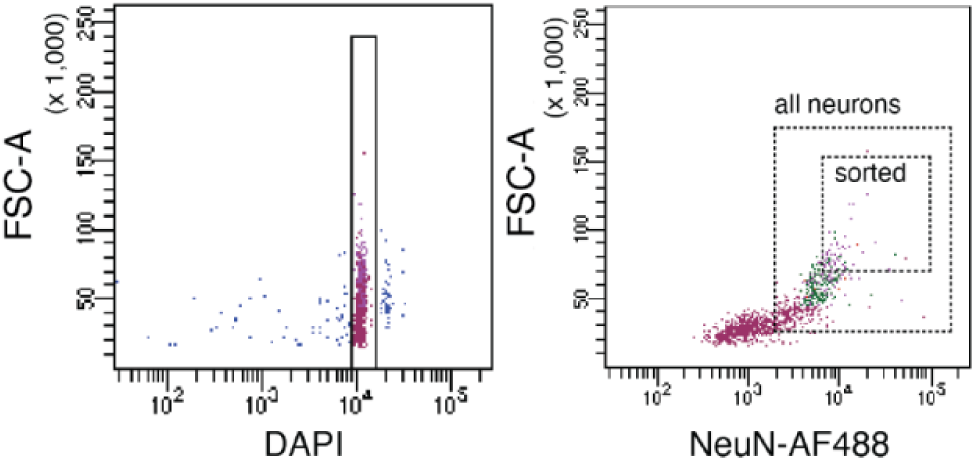
Fluorescence-activated nuclear sorting (FANS) to isolate neurons. (*left*) Staining with 4′,6-diamidino-2-phenylindole (DAPI) is used to identify intact diploid cellular nuclei. (*right*) AF488-conjugated anti-NeuN antibody is used to label large neuronal nuclei for separation from glia and other cell types. The larger box represents all neurons (NeuN+). The smaller box represents the subset of neurons that show high NeuN signal and high forward scatter area (FSC-A) signal, which were sorted for single-cell whole-genome sequencing in this study. Nuclei isolated in this manner represent > 99% pyramidal excitatory neurons (*18*).

**Fig. S2.**
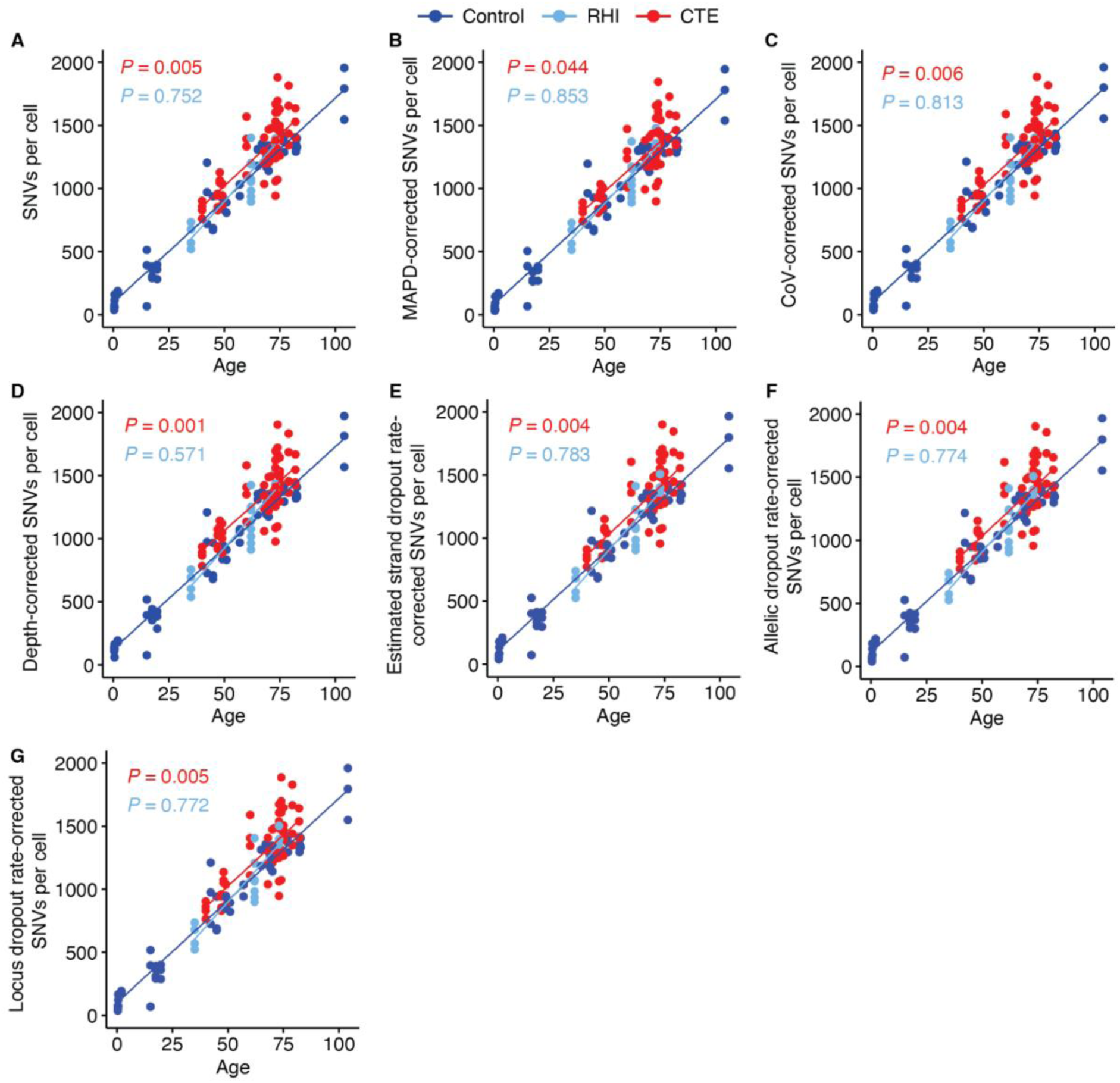
Somatic SNV burden in CTE (red), RHI (light blue), and neurotypical control (dark blue) neurons after controlling for QC metrics. To evaluate the potential confounding effect of sample and sequencing quality on unadjusted somatic SNV burden (**A**, reproduced from Fig. 1C), we calculated the SNV burden corrected for five metrics (**B**–**G**). After controlling for MAPD (B), CoV (C), sequencing depth (D), estimated strand dropout rate (estimated as the square root of allelic dropout rate) (E), allelic dropout rate (F), and locus dropout rate (G), somatic SNV burden in CTE remains significantly increased compared to neurotypical controls while RHI shows similar burden as neurotypical controls.

**Fig. S3.**
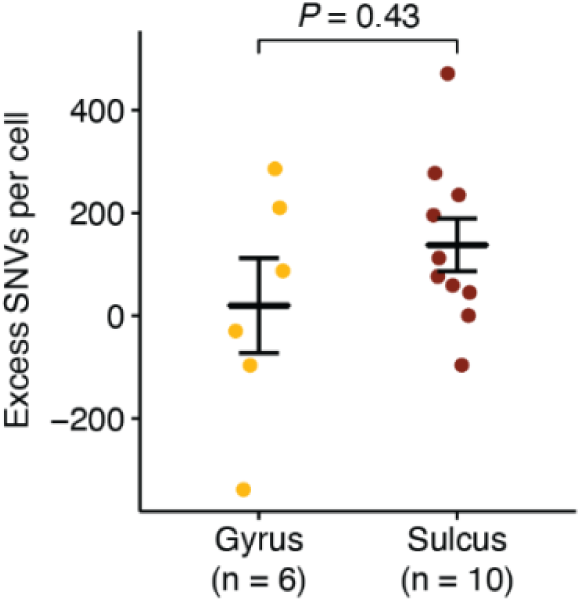
Somatic SNV burden comparison between neurons isolated from gyrus and from sulcus. After normalizing each neuron’s somatic SNV burden by age, there is no significant difference between neurons isolated from gyrus and from sulcus (*P* = 0.43, two-tailed Wilcoxon test). Each point is a single neuron. As we only have three CTE cases with 16 neurons isolated from both loci, the statistical power is rather limited. Data is mean ± standard error.

**Fig. S4.**
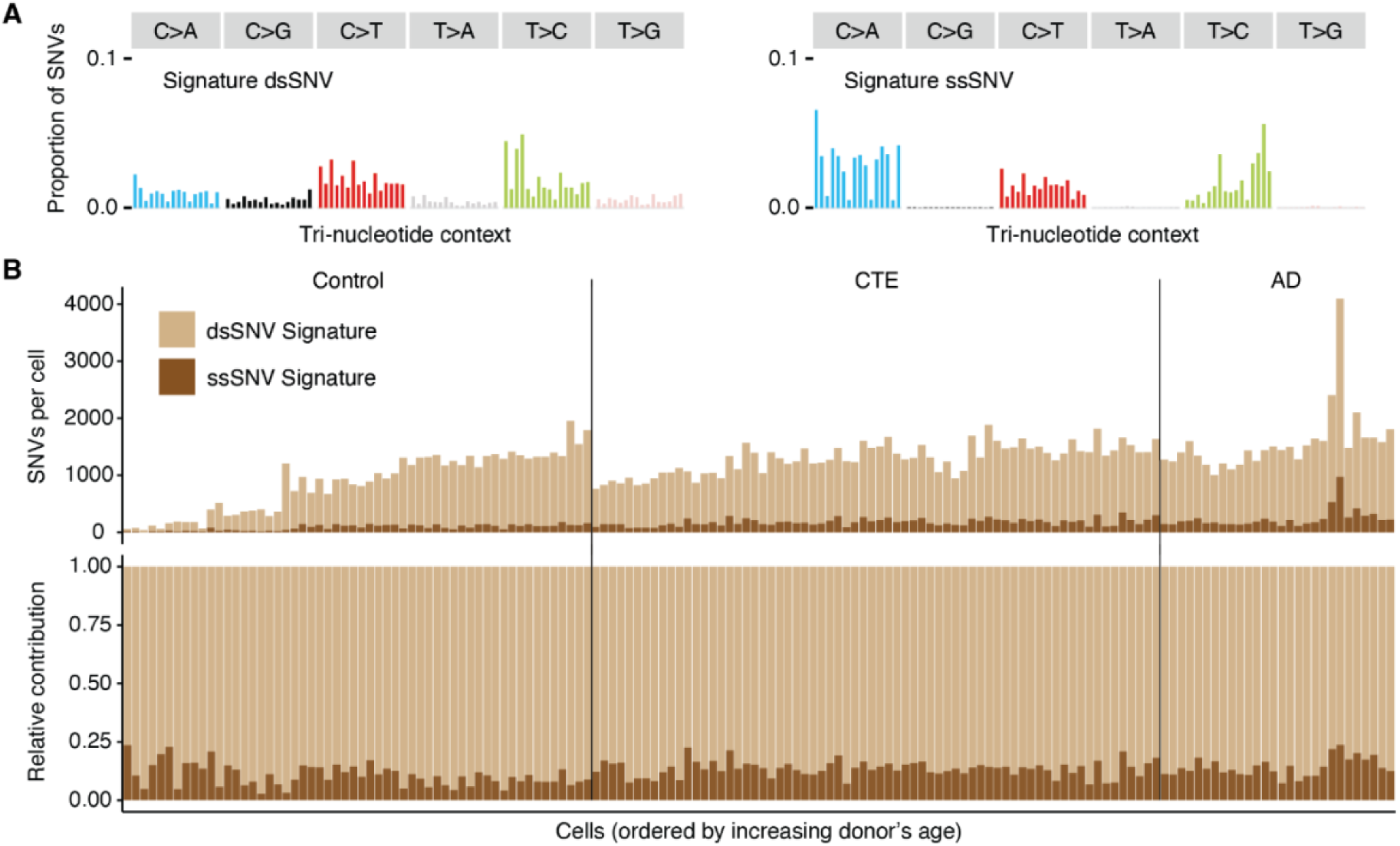
Double-stranded (ds) and single-stranded (ss) SNV signatures extracted from META-CS data and their contribution in PTA neurons across clinical conditions. (**A**) dsSNV and ssSNV signatures are extracted from respective call sets of META-CS data. (**B**) Absolute (*top*) and relative (*bottom*) contribution of dsSNV and ssSNV signatures in PTA-profiled neurotypical control, CTE, and AD neurons where dsSNV shows a predominant contribution (∼87% on average).

**Fig. S5.**
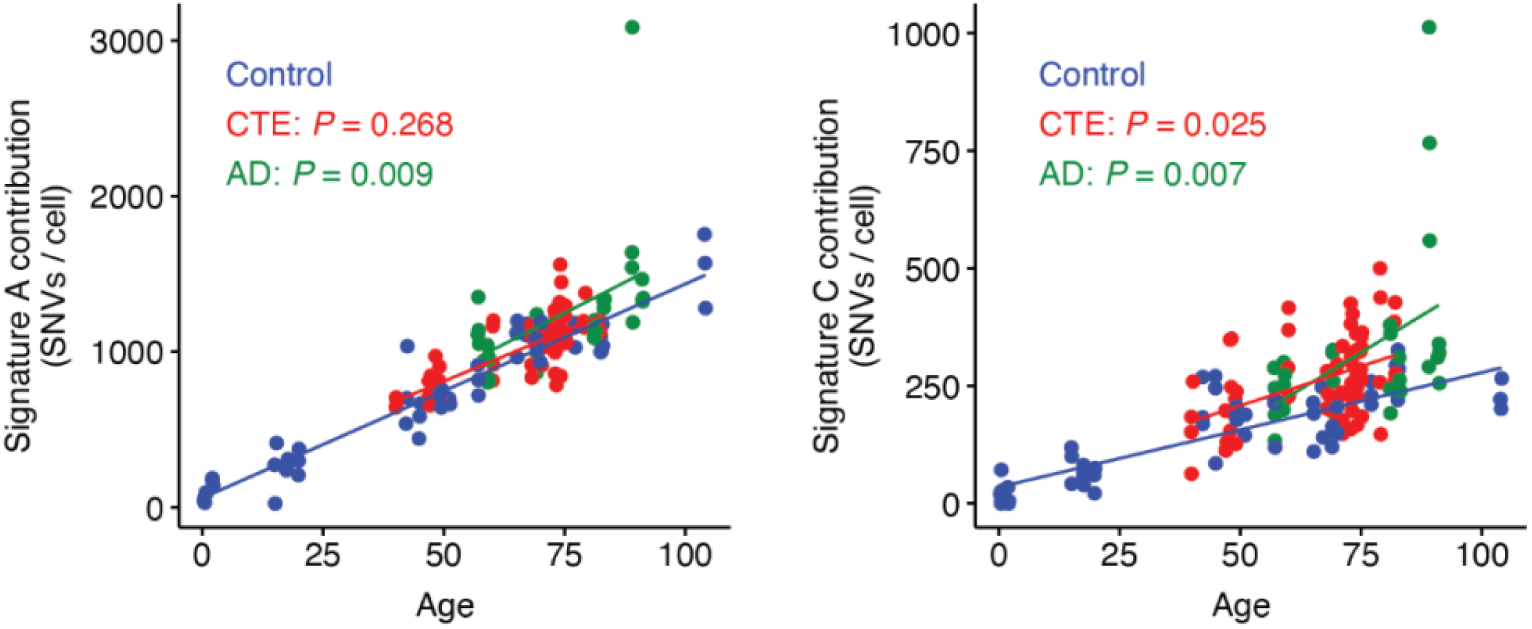
Somatic SNV mutational signatures in CTE, AD, and neurotypical control neurons. Somatic SNV burden separated by contributions from Signatures A (*left*) and C (*right*) in neurotypical control (dark blue), CTE (red), and AD (green) neurons. These signatures decompose somatic SNV burden into age- and disease-specific effects. Their contributions are fitted against age by clinical conditions using a LME model (*left*, Signature A, CTE: *P* = 0.268, AD: *P* = 0.009; *right*, Signature C, CTE: *P* = 0.025, AD: *P* = 0.007).

**Fig. S6.**
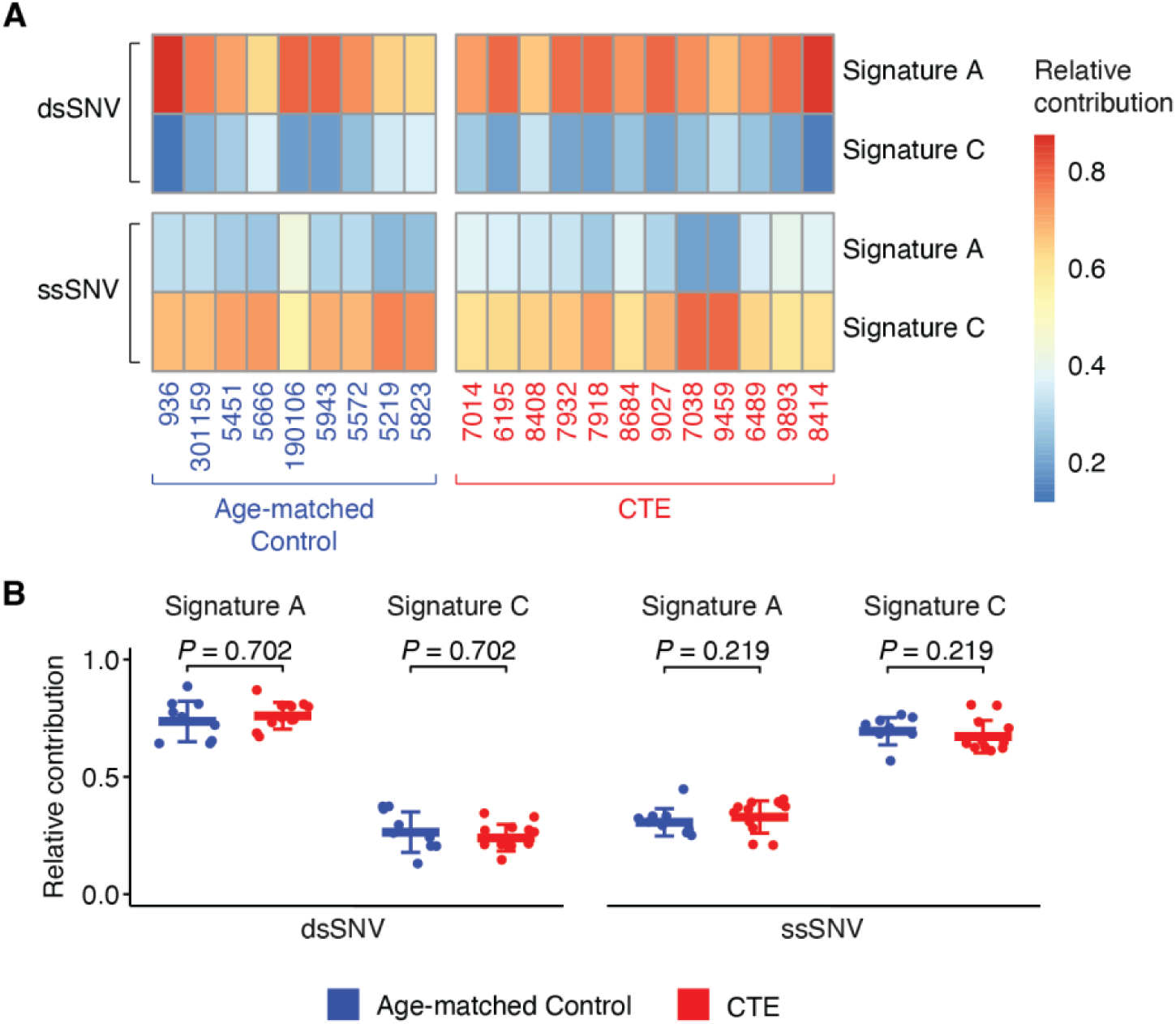
Contribution of Signatures A and C in double-stranded and single-stranded SNVs. (**A**) After decomposing dsSNVs and ssSNVs into Signatures A and C for each case using META-CS data, relative contributions of the two signatures are shown as a heatmap. Case IDs are colored by their group assignment (age-matched control: dark blue, CTE: red). (**B**) Comparison of relative contribution of Signatures A and C to dsSNVs *(left)* and ssSNVs *(right)* between age-matched controls and CTE. Each point represents an individual from META-CS. There is no significant difference between CTE and age-matched controls for either signature. Data is mean ± standard deviation. P-values are from two-tailed Wilcoxon tests.

**Fig. S7.**
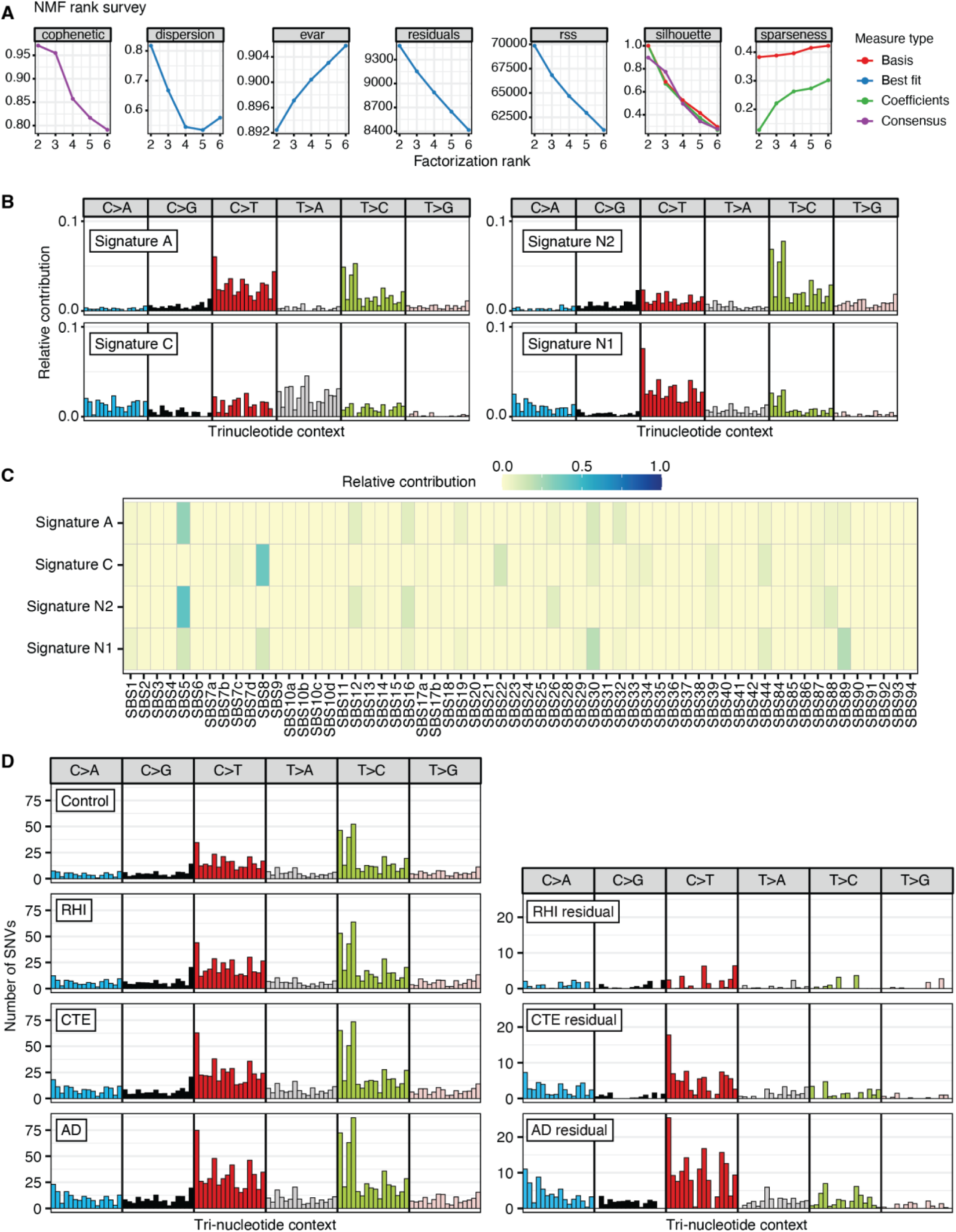
*De novo* signature analysis and mutational patterns of somatic SNVs. (**A**) The optimal number of signatures to extract from PTA data is determined by NMF rank survey after 200 runs and specifically based on the cophenetic coefficient. (**B**) Mutational spectra of two *de novo* signatures N1 and N2 which show a broad resemblance to Signatures C and A, respectively. (**C**) Decomposition of signatures A, C, N1, N2 to COSMIC SBS database. Signature N2 largely resembles Signature A, while Signature N1 carries a mixture of signals from both Signatures A and C, indicating that compared to Signatures A and C, Signatures N1 and N2 mainly capture the same patterns but without a clear distinction in mutational processes. (**D**) Mutational patterns of PTA call sets in neurotypical controls, RHI, CTE, and AD (*left*). After subtracting the pattern of age-matched controls from RHI, CTE, and AD, residual patterns (*right*) are more specific to each clinical condition.

**Fig. S8.**
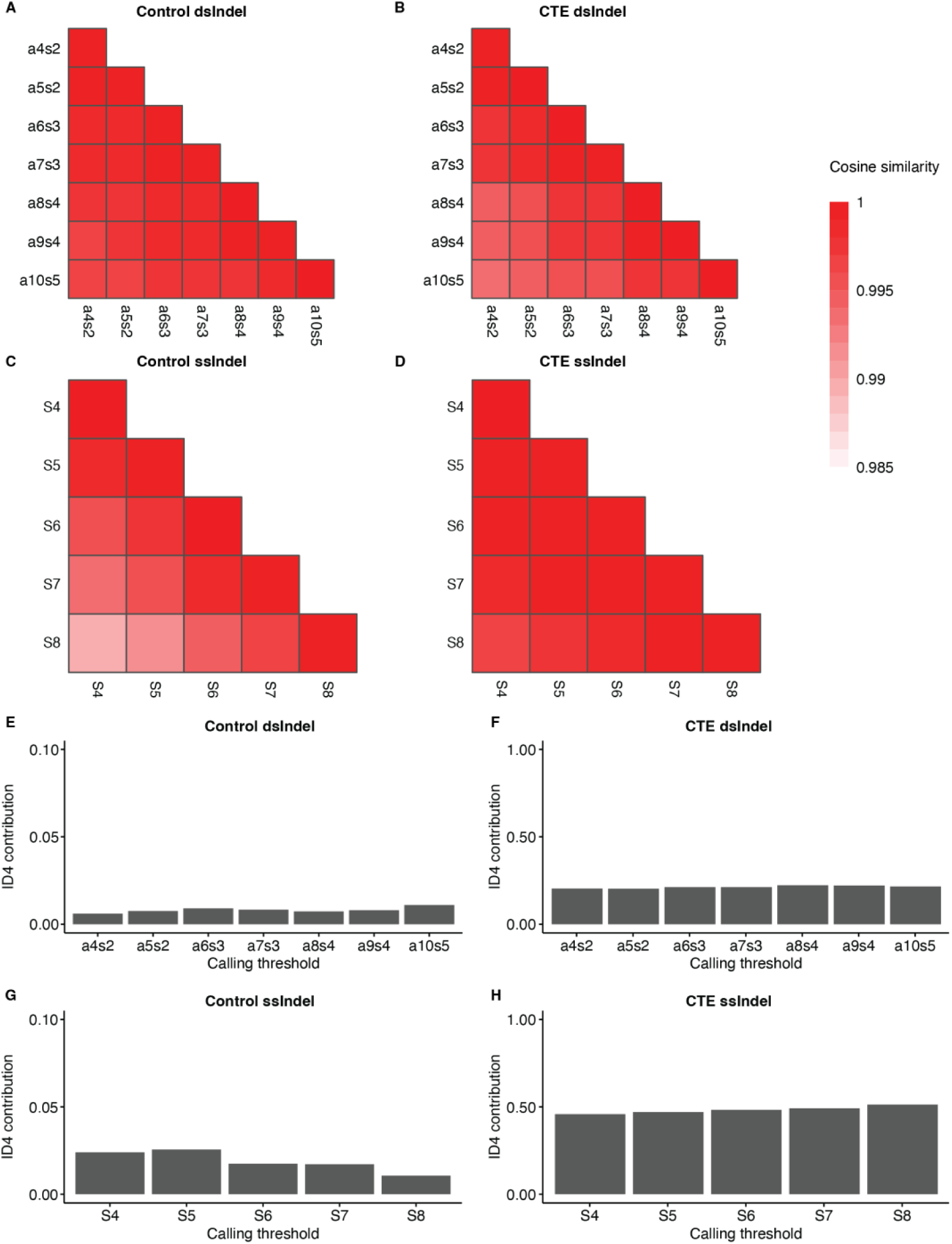
Robustness of META-CS dsIndel and ssIndel signatures and ID4 contribution. (**A**– **D**) Cosine similarity across dsIndel and ssIndel signatures of neurotypical controls and CTE extracted from call sets based on varying calling thresholds. (A, B) Current calling threshold for a dsIndel site is at least 4 total reads (a4) with at least 2 reads from each strand (s2). As calling threshold increases from a4s2 to a10s5, dsIndel signature remains robust in both neurotypical controls and CTE (cosine similarity > 0.98). (C, D) Current calling threshold for a ssIndel site is at least 4 reads from each strand (S4) with REF allele on one strand and ALT allele on the other strand. As calling threshold increases from S4 to S8, ssIndel signature remains robust in both neurotypical controls and CTE (cosine similarity > 0.98). (**E**–**H**) ID4 contribution based on COSMIC decomposition of dsIndel and ssIndel signatures from varying calling thresholds shown in A-D. ID4 contribution is generally consistent even with more stringent calling thresholds.

**Fig. S9.**
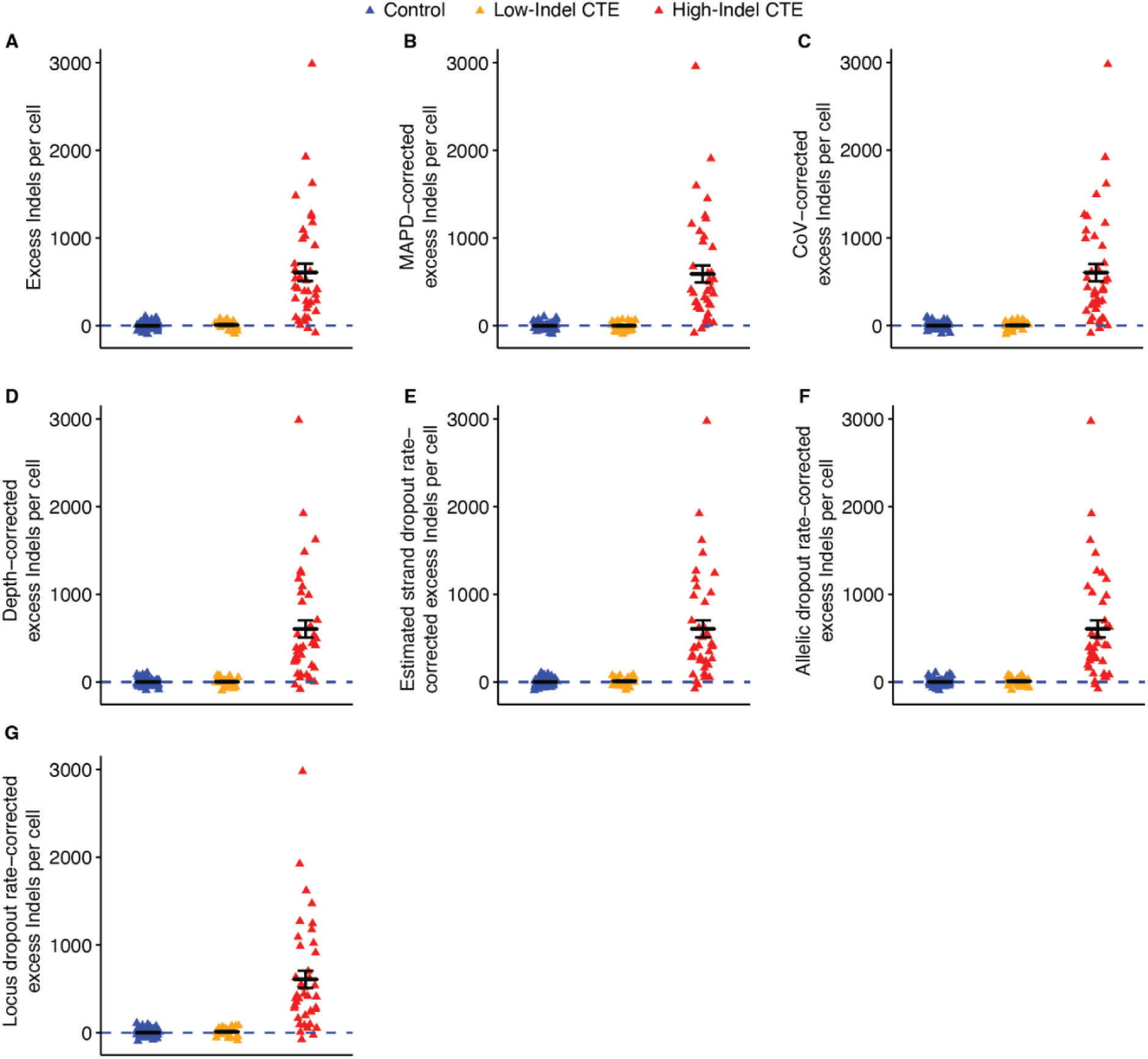
Excess somatic Indels in neurotypical control (dark blue), low-Indel CTE (yellow), and high-Indel CTE (red) after controlling for QC metrics. To evaluate the potential confounding effect of sample and sequencing quality on unadjusted excess somatic Indels (**A**), we calculated the excess Indel burden corrected for five metrics (**B–G**). After controlling for MAPD (B), CoV (C), sequencing depth (D), estimated strand dropout rate (estimated as the square root of allelic dropout rate) (E), allelic dropout rate (F), and locus dropout rate (G), High-Indel CTE consistently has a significant excess of somatic Indels compared to neurotypical controls while Low-Indel CTE shows similar excess Indels as neurotypical controls. Data is mean ± standard error. The dashed blue line shows Indels attributable to age (zero excess).

**Fig. S10.**
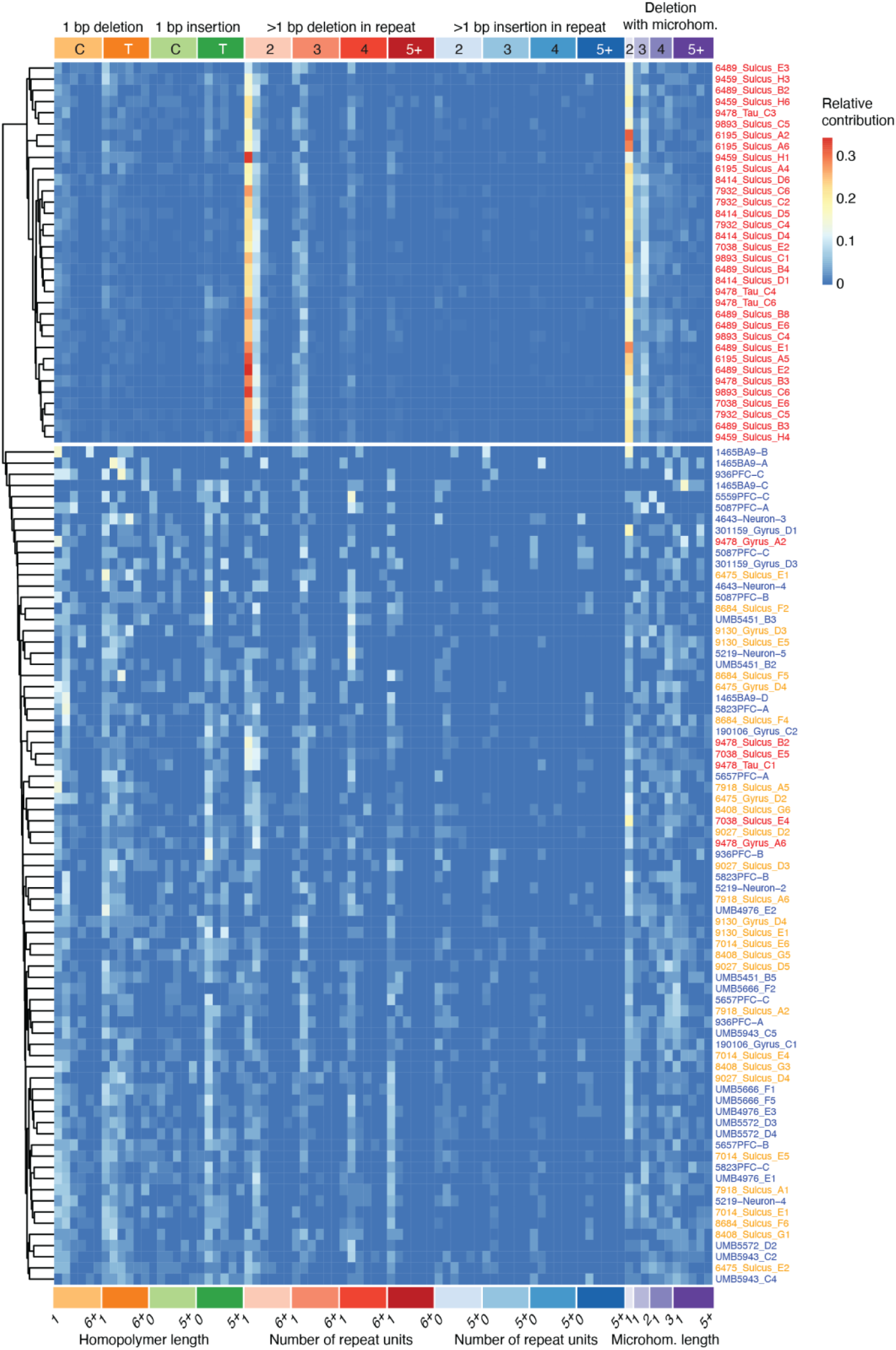
Unsupervised clustering of PTA Indel spectra for each neuron. PTA call sets from single neurons are separated into two clusters using UPGMA hierarchical clustering. Heatmap shows the proportion of 83 Indel types forming the ID83 spectrum on the columns and cells on the rows, where cells from the same individual are colored based on the presence of excess Indels in PTA data (High-Indel CTE: red, Low-Indel CTE: yellow, neurotypical control: blue). Cells from the top cluster represent High-Indel CTE, and cells from the bottom cluster predominantly represent Low-Indel CTE and neurotypical controls.

**Fig. S11.**
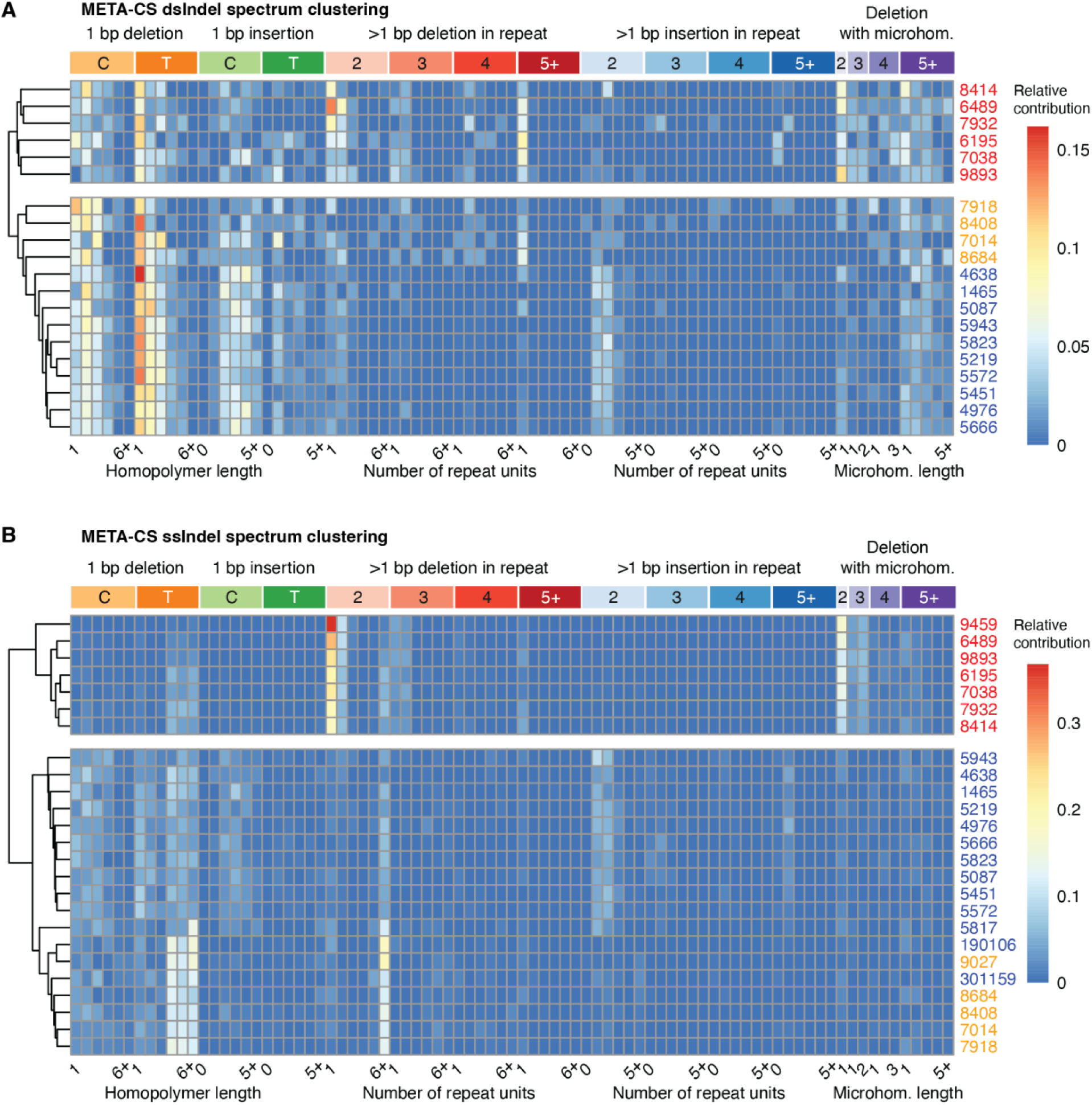
Unsupervised clustering of META-CS dsIndel and ssIndel spectra for each individual. META-CS dsIndel (**A**) and ssIndel (**B**) call sets from single neurons are aggregated to generate individual-specific spectra before separated into two clusters using UPGMA hierarchical clustering. Heatmap shows the proportion of 83 Indel types forming the ID83 spectrum on the columns and individuals on the rows, where individuals are colored based on the presence of excess Indels in their PTA data (High-Indel CTE: red, Low-Indel CTE: yellow, neurotypical control: blue). Clustering results of META-CS are highly consistent with PTA-identified groups in Fig. 4C.

**Fig. S12.**
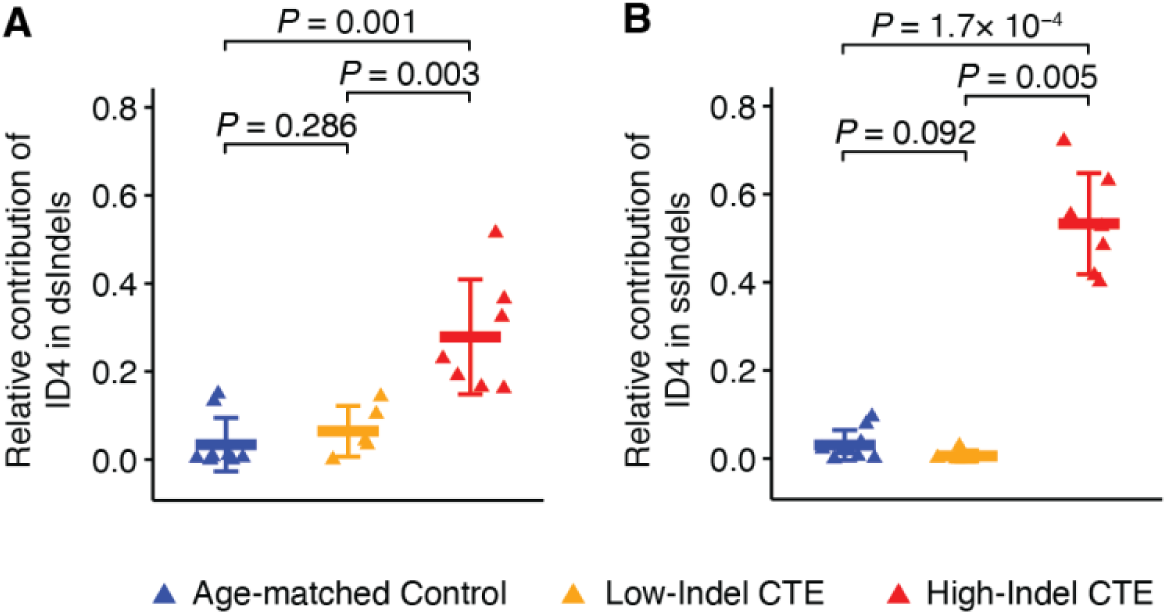
Comparison of relative ID4 contribution to dsIndels and ssIndels across age-matched controls, Low-Indel CTE, and High-Indel CTE. Each triangle represents an individual from META-CS. High-Indel CTE shows a significantly higher ID4 contribution in both dsIndels and ssIndels (A: dsIndel, *P* = 0.001, two-tailed Wilcoxon test; B: ssIndel, *P* = 1.7 × 10^-4^, two-tailed Wilcoxon test), whereas Low-Indel CTE shows no significant difference (A: dsIndel, *P* = 0.286, two-tailed Wilcoxon test; B: ssIndel, *P* = 0.092, two-tailed Wilcoxon test) compared to age-matched controls. Data is mean ± standard deviation.

**Fig. S13.**
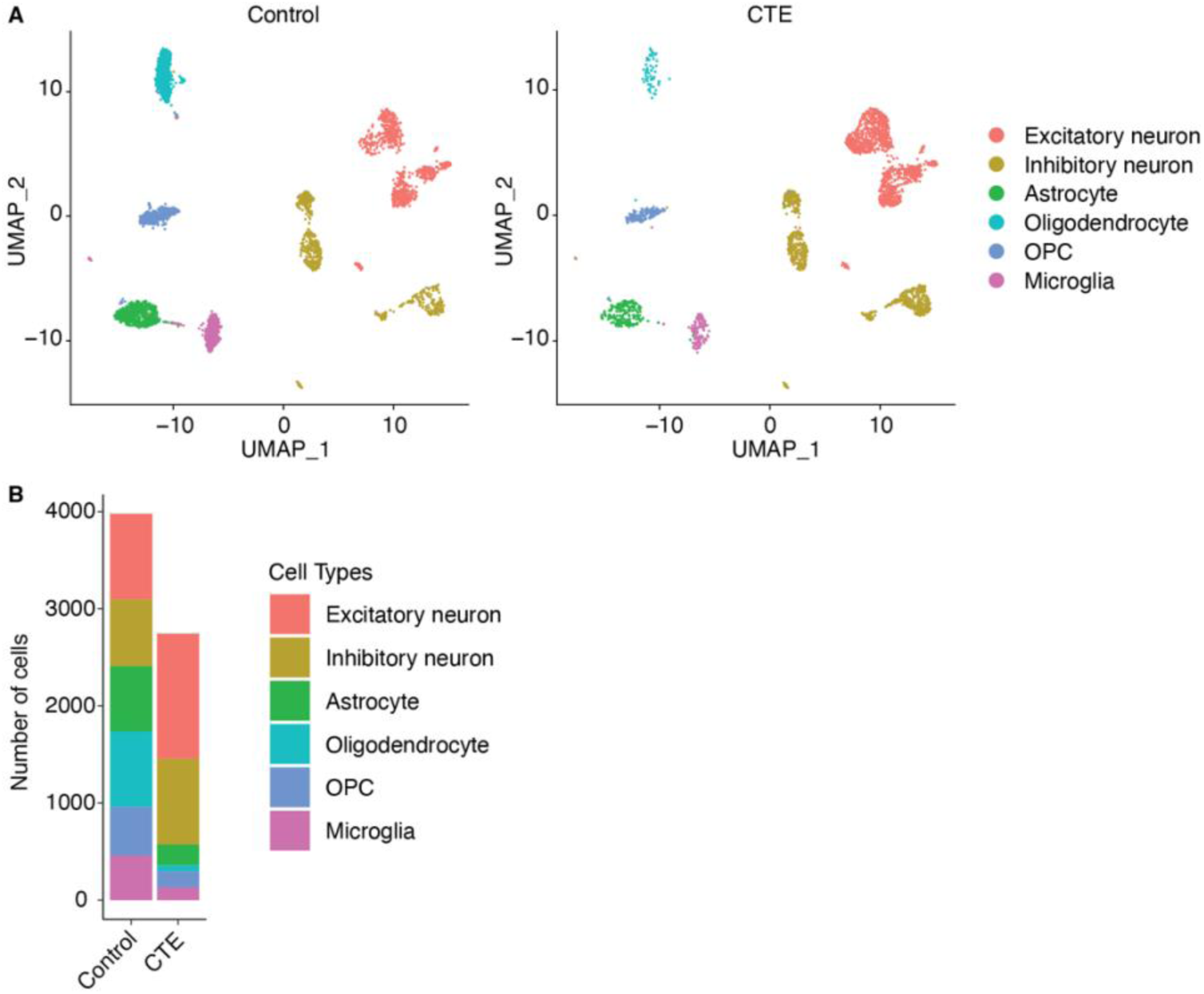
Single-nucleus transcriptomic profiles of different cell types in neurotypical control and CTE samples. (**A**) Profiled cells from neurotypical control (*left*) and CTE (*right*) are plotted based on uniform manifold approximation and projection (UMAP) dimension reduction and color by annotated cell types. (**B**) Number of cells of each cell type in neurotypical control and CTE with a substantial number of neurons in both samples which are used for enrichment analyses.

**Fig. S14.**
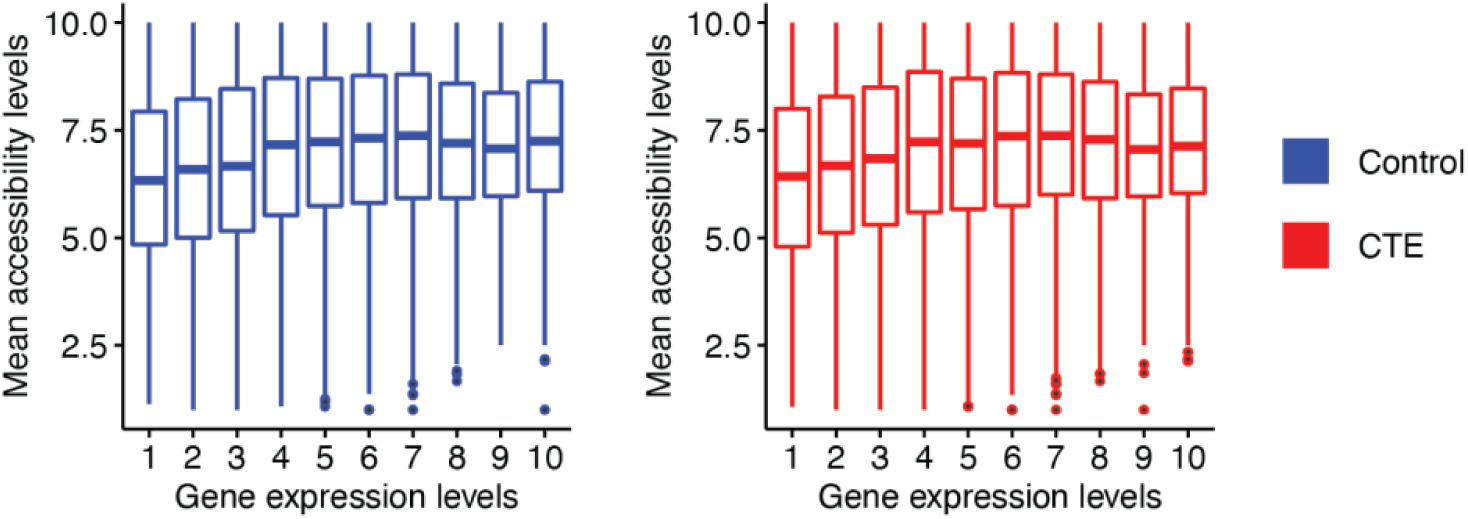
Correlation of gene regions ranked based on gene expression levels and mean accessibility levels. For gene regions assigned to deciles 1 through 10 based on increasing gene expression levels in neurotypical control (*left*) and CTE (*right*), their mean accessibility levels are shown in box plots (bars from top to bottom show the first, second (median), and third quartile; whiskers extend 1.5 IQR with outlier data points shown separately).

**Table S1.**

Sample information. Clinical data, library and sequencing metrics.

**Table S2.**

PTA and META-CS sequencing statistics.

**Table S3.**

PTA SNV and Indel rates.

**Table S4.**

PTA SNV and Indel calls.

**Table S5.**

META-CS SNV and Indel calls.

**Table S6.**

Gene Ontology terms enriched for SNVs and Indels.

